# Genetic deletion of genes in the cerebellar rhombic lip lineage can stimulate compensation through adaptive reprogramming of ventricular zone-derived progenitors

**DOI:** 10.1101/383778

**Authors:** Alexandre Wojcinski, Morgane Morabito, Andrew K. Lawton, Daniel N. Stephen, Alexandra L. Joyner

## Abstract

**Background:** The cerebellum is a foliated posterior brain structure involved in coordination of motor movements and cognition. The cerebellum undergoes rapid growth postnataly due to Sonic Hedgehog (SHH) signaling-dependent proliferation of ATOH1+ granule cell precursors (GCPs) in the external granule cell layer (EGL), a key step for generating cerebellar foliation and the correct number of granule cells. Due to its late development, the cerebellum is particularly vulnerable to injury from preterm birth and stress around birth. We recently uncovered an intrinsic capacity of the developing cerebellum to replenish ablated GCPs via adaptive reprogramming of Nestin-expressing progenitors (NEPs). However, whether this compensation mechanism occurs in mouse mutants affecting the developing cerebellum and could lead to mis-interpretation of phenotypes was not known.

**Methods:** We used two different approaches to remove the main SHH signaling activator GLI2 in GCPs: 1) our mosaic mutant analysis with spatial and temporal control of recombination (MASTR) technique to delete *Gli2* in a small subset of GCPs; 2) An *Atohl-Cre* transgene to delete *Gli2* in most of the EGL. Genetic Inducible Fate Mapping (GIFM) and live imaging were used to analyze the behavior of NEPs after *Gli2* deletion.

**Results:** Mosaic analysis demonstrated that SHH-GLI2 signaling is critical for generating the correct pool of granule cells by maintaining GCPs in an undifferentiated proliferative state and promoting their survival. Despite this, inactivation of *GLI2* in a large proportion of GCPs in the embryo did not lead to the expected dramatic reduction in the size of the adult cerebellum. GIFM uncovered that NEPs do indeed replenish GCPs in *Gli2* conditional mutants, and then expand and partially restore the production of granule cells. Furthermore, the SHH signaling-dependent NEP compensation requires *Gli2*, demonstrating that the activator side of the pathway is involved.

**Conclusion:** We demonstrate that a mouse conditional mutation that results in loss of SHH signaling in GCPs is not sufficient to induce long term severe cerebellum hypoplasia. The ability of the neonatal cerebellum to regenerate after loss of cells via a response by NEPs must therefore be considered when interpreting the phenotypes of conditional mutants affecting GCPs.

## Background

The cerebellum (CB) consists of 80% of the neurons in the human brain (1) (60% in mouse (2)), and is involved in balance and motor coordination, but also modulates language, reasoning and social processes via its connections throughout the forebrain (3-7). The CB undergoes its major growth in the third trimester and infant stage in humans, and the first 2 weeks after birth in mice, primarily due to expansion of the granule cell precursor (GCP) pool in the external granule cell layer (EGL) (8-10). Given the late development of the CB compared to other brain regions, the CB is particularly sensitive to environmental and clinical factors that impact on growth (or cause injury) around birth. Furthermore, CB hypoplasia and prenatal injury is the second leading factor associated with autism (11). It is therefore important to identify genes that regulate cerebellum development. Many of the genes have been identified based on motor defects in homozygous null mutant mice, or in conditional mutants that remove genes in specific cell lineages. Intrinsic growth compensation mechanisms involving lineages where the gene does not function could however, obscure the normal function of a gene in cerebellar growth.

The CB develops from two germinal zones. The ventricular zone (VZ) gives rise to all the inhibitory neurons, including Purkinje cells (PCs) (12) as well as Nestin-expressing progenitors (NEPs) that expand in the cerebellar cortex after birth to produce astrocytes, including specialized Bergmann glia, and late born interneurons of the molecular layer (13, 14). *Ptfla^Cre^* mice have been used to delete genes in inhibitory neurons and some glia (15). Excitatory neurons including granule cells (GCs) originate from the upper rhombic lip (16-18). In mice, the EGL is established between embryonic day (E) 13.5 and

E15.5. *Atoh1*-expressing GCPs then proliferate and expand in the EGL until ~postnatal day (P) 15 in response to Sonic Hedgehog (SHH) secreted by PCs (19-21). When GCPs become postmitotic, they migrate down Bergmann glial fibers to form the internal granule cell layer (IGL). Interestingly, in rodent models the developing CB has been found to have a remarkable ability to recover from some injuries (22-24). Indeed, we recently found that proliferating cerebellar GCPs can be replaced via adaptive reprogramming of NEPs after an acute depletion of the perinatal EGL due to irradiation (25-27). Thus, NEPs in the neonatal CB have highly plastic behaviors. However, whether NEPs are harnessed to replenish cells lost in developmental mutants that lack key factors required for expansion and survival of GCPs has not been addressed.

One of the major pathways driving CB development is HH signaling. There are three hedgehog *(Hh)* genes in mammals, *Indian* (Ihh), *Desert (Dhh)* and *Shh* (28, 29). *Shh*, the most widely expressed *Hh* gene, is required for development of most organs (29) by regulating a variety of cell behaviors including cell death, proliferation, specification and axon guidance. The cellular context (i.e. tissue, developmental stage, convergence of other signaling pathways) and concentration of SHH are thought to determine the particular response of a cell to SHH. HH signal transduction is mediated by the receptors Patched1 (PTCH1) and Smoothened (SMO) (28-30). In the absence of HH signaling, PTCH1 constitutively represses SMO activity, whereas HH binding relieves this inhibition, in part by allowing accumulation of SMO in cilia (31). The GLI/Ci transcription factors are the effectors of the HH pathway. In mammals, the transcriptional activator (A) and repressor (R) functions of the GLIs have been divided between the three proteins (32). A general rule is that high levels of HH signaling induce the formation of a GLI2 activator (GLI2^A^) and this leads to transcription and translation of an addition activator, GLI1^A^, while a reduction or absence of the ligand allows for the formation of a GLI3 repressor (GLI3^R^). Importantly, we demonstrated that *Gli1* expression is dependent on GLI2/3^A^, and thus is only expressed in cells receiving a high level of HH signaling (33, 34). The three *Gli* genes, *Shh, Smo, Ptch1* and *Ptch2* are expressed in the CB and all but *Ptch2* are required for CB development (20, 21, 35-37). In particular, we have shown that SHH functions by inducing GLI1^A^/2^A^ and is required for expansion of GCPs, primarily after birth (20, 38), whereas Gli3 is not required in the cerebellum after E12.5 (36). In addition to the crucial role of SHH in generating the pool of GCs, expansion of NEPs and thus production of NEP-derived interneurons and astroglia (astrocytes and Bergmann glia) also require SHH-signaling ((13, 25, 39)). Furthermore, HH-signaling in NEPs is crucial for expansion of NEPs, recovery of the EGL and scaling of interneuron numbers after injury to the EGL (25).

Here we report that deletion of *Gli2* in the vast majority of the GCPs is not sufficient to induce major cerebellar hypoplasia. Using our MASTR technique (40) in a mosaic mutant analysis of the effect of deleting *Gli2* in scattered GCPs, we found that HH/GLI2-signaling is indeed necessary to maintain GCPs in an undifferentiated and proliferative state and to promote their survival. However, and similar to when the EGL is depleted using irradiation, we uncovered that NEPs are harnessed to repopulate the EGL and then wild type progenitors differentiate into GCs when *Gli2* is deleted in most GCPs using an Atoh1-driven constitutive Cre (41). Our results not only provide more evidence for the unusual ability of the CB to recover from perinatal stress, but also reveal that NEP-dependent compensation should be taken into account when studying genes implicated in GCP development and when using the *Atoh1-Cre* transgene.

## Methods

### Animals

The following mouse lines were used: *Gli2^flox/flox^* (20), *Atoh1-Cre* (41), *Atoh-FlpoER, Nestin-FlpoER* (a transgene similar to that described in (40)) and *Rosa26^MASTR(frt-STOP-frt-GFPcre^*) (40), *Atoh1-GFP* (42), *Nes-CFP* (43), *Rosa26^FRT-STOP-FRT-TDTom^* (*Jackson Laboratory, 021875*). The *Atoh-FlpoER* line, was made using the FLPoER1 cDNA described in (40) by subcloning it into the *Atoh1* expression construct described in (17). All mouse lines were maintained on an outbred Swiss Webster background and both sexes were used for the analysis. Animals were housed on a 12 h light/dark cycle and were given access to food and water *ad libitum*. All experiments were performed using mice ages P0–P30.

Tamoxifen (Tm, Sigma-Aldrich) was dissolved in corn oil (Sigma-Aldrich) at 20mg/ml. P2 *Atoh1-FlpoER*/+; *R26*^*MASTR*/+^; *Gli2*^*flox/flox*^, *Atoh1-FlpoER*/+; *R26*^*MASTR*/+^; *Gli2*^*flox/flox*^ and P0 *Nes-FlpoER*/+; *R26*^*FSF-TDTom*/+^, *Nes-FlpoER*/+; *R26*^*FSF-TDTom*/+^; *Atoh-GFP*/+ mice as well as *Nestin-FlpoER*/+; *R26*^MASTR/+^; *Gli2^flox/flox^*, *Atoh1-Cre*/+; *Gli2^flox/flox^* and *Nestin-FlpoER*; *R26*^*MASTR*/+^; *Atoh1-Cre*/+; *Gli2^flox/flox^* mice and littermate controls received one 200μg/g dose of Tm via subcutaneous injection.

50 μg/g 5-ethynyl-2_-deoxyuridine (EdU; Invitrogen) was administered via intraperitoneal injection (10mg/ml in sterile saline) one hour before the animals were sacrificed.

### Tissue Processing, immunohistochemistry (IHC) and transcript detection

For animals younger than P4, they were anaesthetized by cooling and brains were dissected out and fixed in 4% paraformaldehyde overnight at 4°C. Animals P4-30 received 50 μl intraperitoneal injections of ketamine and received ice-cold PBS via transcardial perfusion followed by 4% paraformaldehyde. Brains were collected and submersion fixed in 4% paraformaldehyde overnight at 4°C. Tissues were processed for frozen embedding in optimal cutting temperature (OCT) compound and sectioned in the parasagittal plane on a Leica cryostat at 12 μm. For IHC, sections were incubated overnight at 4°C with the following primary antibodies: rabbit anti-Ki67 (Thermo Scientific, RM-9106-S0), mouse anti-P27 (BD Pharmigen, 610241), rabbit anti-PAX6 (Millipore, AB2237), goat anti-GLI2 (R&D System, AF3635), Goat anti-SOX2 (R&D System, AF2018), rabbit anti-GFP (Life Technologies, A11122), rat anti-GFP (Nacalai Tesque, 04404-84), mouse anti-NeuN (Millipore, MAB377) diluted in PBS with 5% BSA (Sigma-Aldrich) and 0.3% Triton X-100 (Fisher Scientific). Sections were then exposed for 2h at room temperature to secondary species-specific antibodies conjugated with the appropriate Alexa Fluor (1:500; Invitrogen). EdU was detected using a commercial kit (Life Technologies) after the IHC reactions. TUNEL staining and *in situ* hybridization were performed according to standard protocols. *Cre* and *Gli1* cDNAs were used as the template for synthesizing digoxygenin-labeled riboprobes. Images were collected on a DM6000 Leica microscope and processed using Photoshop software.

### Live imaging

*Ex vivo* cerebellar slice culture was done as previously described (25). Briefly, P8 cerebella were embedded in 2.5% low-melting point agarose and saggitally sliced at 250μM on a Vibratome. Slices were immediately taken to either a Leica TCS SP8 or SP5 confocal microscope platform. Slices were maintained in Eagle’s Basal Medium with 2mM L-glutamine, 0.5% glucose, 50U/ml Penicillin-streptomycin, 1xB27 and 1xN2 supplements at 37°C and 5% CO_2_. Image stacks were acquired every 5min for ~4h. Cell tracking was performed using Imaris software. The autoregressive tracking function was employed with a spot size of 6μM and a step size of 7μM. Manual correction was performed.

### Quantifications and Statistical Analyses

ImageJ software was used to measure the area (mm^2^) of cerebellar sections near the midline. For all IHC staining, cell counts were obtained using ImageJ and Neurolucida Software. For each developmental stage, three sections were analyzed per animal and ≥3 animals. Statistical analyses were performed using Prism software (GraphPad) and significance was determined at *P* < 0.05. All statistical analyses were two-tailed. For two-group comparisons with equal variance as determined by the *F*-test, an unpaired Student’s *t* test was used. Welch’s correction was used for unpaired *t*-tests of normally distributed data with unequal variance. P values are indicated in the figures. No statistical methods were used to predetermine the sample size, but our sample sizes are similar to those generally employed in the field. No randomization was used. Data collection and analysis were not performed blind to the conditions of the experiments.

## Results

### Mosaic analysis reveals SHH-GLI2 signaling is critical for maintaining GCPs in an undifferentiated proliferative state and promoting their survival

Our previous studies demonstrated that loss of the majority of HH-signaling in the entire CB at mid-gestation (*Nes-Gli2* conditional knockout or CKO - *Nestin-Cre; Gli2^flox/flox^* mice) results in an almost complete lack of GCPs at birth and a very diminished CB in adults (20). Since HH-signaling is required after birth in NEPs for their expansion and production of late born interneurons and astrocytes in the CB (13), it is possible that part of the phenotype observed in *Nes-Gli2* CKOs was due to loss of HH-signaling in Non-GCP cells. We therefore took two approaches to test the cell autonomous requirement for HH signaling in GCPs. First we used the *R26^FSF-GFPcre^* MASTR allele (*R26^MASTR^*) (40) and a *Atoh1-FlpoER* transgene to knock out *Gli2* in scattered GCPs at ~P3 by administering Tamoxifen (Tm) at P2 and analyzed the percentage of undifferentiated GFP+ GCPs (GFP+ cells in the proliferating outer EGL/total GFP+ cells – proliferating and post mitotic) at both P4 and P8 (Fig. 1a-c). We did indeed observe a significant decrease in the percentage of GFP+ cells that were GCPs in the medial CB (vermis) of P8 *Atoh1-M-Gli2* CKOs (*R26*^*MASTR*/+^; *Atoh1-FlpoER*/+; *Gli2*^*flox/flox*^ mice; n=3;) compared to *Atoh1-M-Gli2* heterozgous (het) controls (*R26*^MASTR^/+; *Atoh1-FlpoER*/+; *Gli2*^*flox*/+^ mice; n=3) (29.79% compared to 67.09%) (Fig. 1d). Using a 1hr pulse of EdU, we found that the proliferation index (#EdU+ GFP+ cells in the outer EGL/total GFP+ cells in the outer EGL) of *Atoh1-M-Gli2* CKO GCPs was significantly decreased compared to controls (n=3; 14.39% compared to 29.06%) (Fig. 1e). At P4 however, there was no significant difference in the percentage of undifferentiated GFP+ GCPs between *Atoh1-M-Gli2* CKOs and controls (p=0.162) (Fig. 1d). Although not significant, we observed a trend towards a decrease in the proliferation index in P4 *Atoh1-M-Gli2* CKO cerebella (CKO vs control, p=0.162) (Fig. 1e). Interestingly, at P4, only 2 days after Tm injection, the number of GFP+ cells in the oEGL already appeared decreased in mutants compared to controls (CKO vs control, p=0.081) (Fig. S1a) suggesting that some of the cells underwent cell death. Consistent with this idea, TUNEL staining revealed a significant increase in cell death in the entire EGL at P4 (69.44±7.76/section in mutants vs 37.67±5.1 in controls, p=0.027). We performed the same analysis in the lateral CB (hemispheres) and found a similar outcome (Fig. S1b-d). These results reveal that HH-signaling through GLI2 plays an important role in maintaining GCPs in an undifferentiated state, and also promotes their proliferation and survival.

**Fig. 1.**
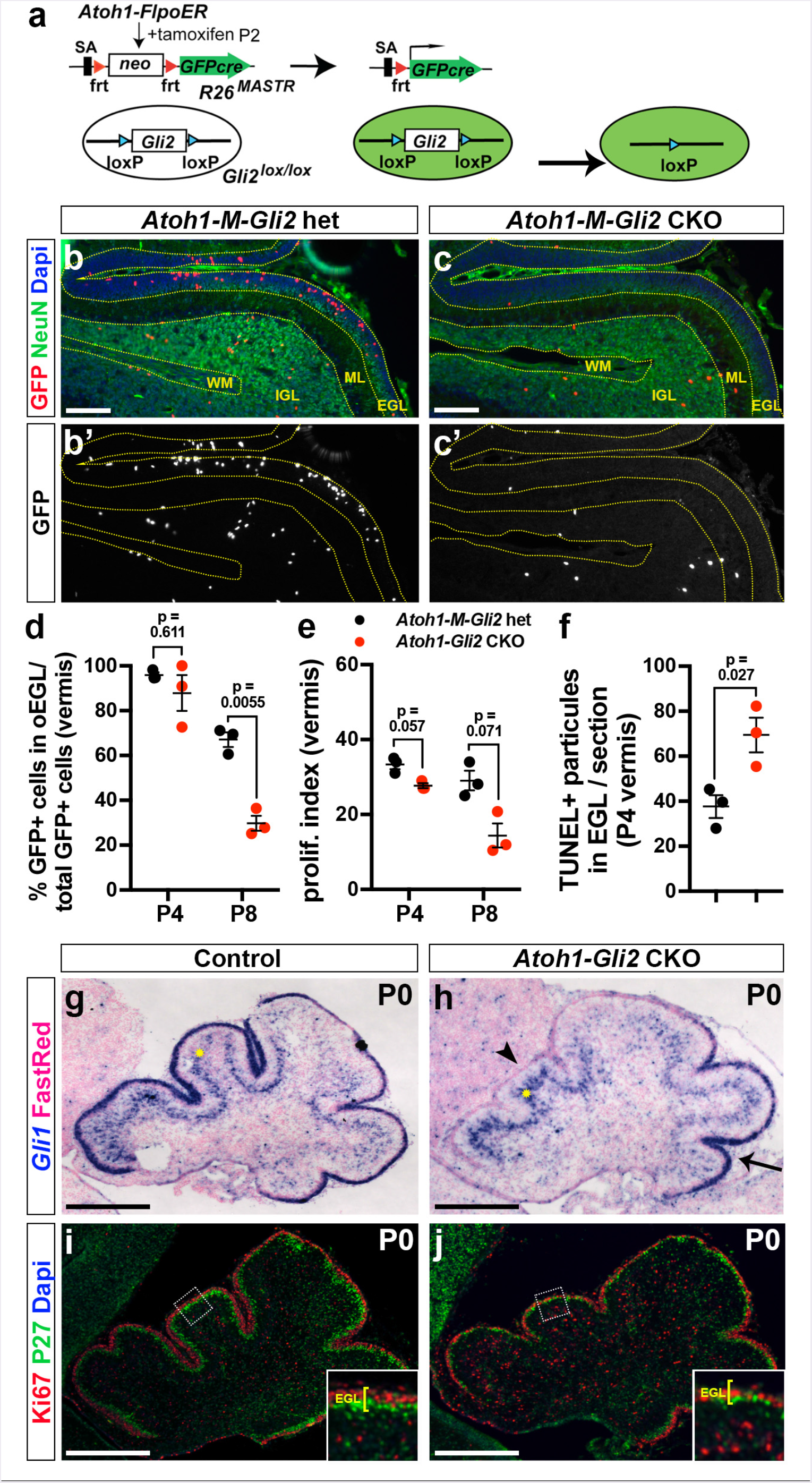
HH-GLI2 signaling maintains GCP in an undifferentiated state and promotes their survival. (**a**) Schematic representation of the MASTR approach. The *R26*^*MASTR*^ allele expresses a GFPcre fusion upon Flp induced deletion of a *neo* (STOP) cassette. When the *R26^mastR^* allele and the *Atoh1-FlpoER* transgene are combined with a floxed gene such as *Gli2*, recombination of the loxP sites occurs in >98% of GFP+ cells within 3 days of administrating tamoxifen (Tm) at P2. The mutant cells and their progeny can subsequently be identified by the continuous expression of eGFP from the *R26* allele. (**b-c**) Fluorescent Immuno-Histo-Chemical (FIHC) detection of the indicated proteins and dapi on mid-sagittal sections (lobule VII and VIII) of P8 control *R26*^*MASTR*/+^; *Atoh1-FlpoER*/+; *Gli2*^*lox*/+^ (*Atoh1-M-Gli2* het, **b**) and *R26*^*MASTR*+^; *Atoh1-FlpoER*/+; *Gli2^lox/lox^* (*Atoh1-M-Gli2* CKO, **c**) mice treated Tm at P2. (**d-f**) Graphs of the proportion of GFP+ cells in the outer (o) EGL at P8 (n=3) (**d**), the proliferation index at P8 (% [GFP+ EdU+] cells of all [GFP+] cells in the oEGL) (n=3) (**e**) and the number TUNEL+ particles per section at P4 (n=3) (**f**) in *Atoh1-M-Gli2* het (control, black) and *R26*^*MASTR*/+^; *Atoh1-FlpoER*/+; *Gli2^lox/lox^* (*Atoh1-M-Gli2* CKO, red) mice treated Tm at P2. All of the analyses were performed on 3 midline sections per brain. All graphical data are presented as means ± SEM and significance determined using two-tailed T-test. (**g-h**) *In situ* hybridization of *Gli1* mRNA on P0 mid-sagittal cerebellar sections of *Gli2^lox/lox^* (control,**g**) and *Atoh1-Cre*/+; *Gli2^lox/lox^* (*Atoh1-Gli2* CKO, **h**). Arrowheads indicate the loos of *Gli1* expression in the EGL and yellow asterisks indicate *Gli1* expression in Bergmann glia in the Purkinje Cell Layer (PCL). (**i-j**) FIHC detection of the indicated proteins and dapi on P0 mid-sagittal cerebellar sections of *Gli2^lox/lox^* (control, **I**) and *Atoh1-Cre*/+; *Gli2^lox/lox^* (*Atoh1-Gli2* CKO, **J**). High power images are shown of the areas indicated by white rectangles and the thickness of the EGL is indicated by yellow bracket. Scale bars represent 100μm (**b-c**) and 500μm (**g-j**).

As an alternative approach to a mosaic mutant analysis, we deleted *Gli2* in the vast majority of GCPs (*Atoh1-Cre*/+; *Gli2*^*flox/flox*^ or *Atoh1-Gli2* CKOs). Consistent with previous studies using whole cerebellum *Cre* transgenes and our mosaic analysis, at P0 the anterior vermis of *Atoh1-Gli2* CKOs (n=5) was consistently smaller than controls *(Gli2^flox/flox^)* and the EGL was greatly diminished (Fig. 1g-j). SHH-GLI2 signaling loss was confirmed by the lack of *Gli1* expression in the EGL of *Atoh1-Gli2* CKO cerebella (Fig. 1G-H). Moreover, proliferation (Ki67) and differentiation (P27) were disrupted in the mutant EGL since two distinct EGL layers were not present (Fig. 1i-j). Interestingly, we observed an apparent increase of *Gli1* expression in the Purkinje cell layer (PCL) suggesting that deletion of *Gli2* in the EGL induced a cell non-autonomous up-regulation of HH-signaling in the Bergmann glia in the PCL (star in Fig. 1g-h). The lack of a phenotype in the posterior vermis is likely explained by low expression of *Cre* (44) in this region and thus low recombination (45) (Fig.S2).

All together, these results confirm a major role played by SHH-signaling through GLI2 to promote the expansion of the EGL and thus ensure the generation of the correct number of GCs.

### The size of the *Atoh1-Gli2* CKO cerebellum progressively recovers after birth

We have recently shown that the size of the CB can recover to ~80% its normal size after postnatal injury (irradiation) to the EGL (25). To test whether genetic ablation of *Gli2* in the EGL can trigger a similar recovery mechanism, we analyzed the phenotype of adult (P30) *Atoh1-Gli2* CKO cerebella. The area of mid-sagittal sections of P30 animals was quantified, and revealed only a 21.7 ± 12.0% reduction (n=6) in *Atoh-Gli2* CKOs compared to littermate controls (Fig. 2a-d). Interestingly, we observed a large variability in the phenotype, with some mutant cerebella recovering better than others and some mice had a cytoachitecture that was very similar to controls (compare Fig.2b and c). We then measured the area of midsagittal cerebellar sections from P4, P8 and P12 mice to determine how recovery occurred in some mice (Fig. 2e-j). As predicted, the size of *Atoh1-Gli2* CKOs cerebella was greatly reduced at P4 compared to control animals (57% the size of control CB) (Fig. 2k-l). and then partially recovered. Since our mosaic mutant results showed a similar behavior of *Gli2* CKO GCPs in the hemispheres and vermis, we analyzed the phenotype of *Atoh1-Gli2* CKOs in the hemisphere. Curiously, unlike the vermis we did not observe a significant decrease in the size of the mutant hemispheres at P30 compared to controls (p=0.152) (Fig. S3a-c). Analysis of hemispheric sagittal sections from P4 (n=3), P8 (n=3) and P12 (n=3) animals revealed the hemispheres were greatly diminished at P4 and then progressively recovered both their size and cytoarchitecture (Fig. S3d-i). Interestingly, whereas the vermis of *Atoh1-Gli2* CKOs mice at P8 showed a clear hypoplasia phenotype, the hemispheres exhibited extra folia (arrow in FigS3), suggesting different compensation responses in the two locations of the CB in *Atoh1-Gli2* CKOs.

**Fig. 2.**
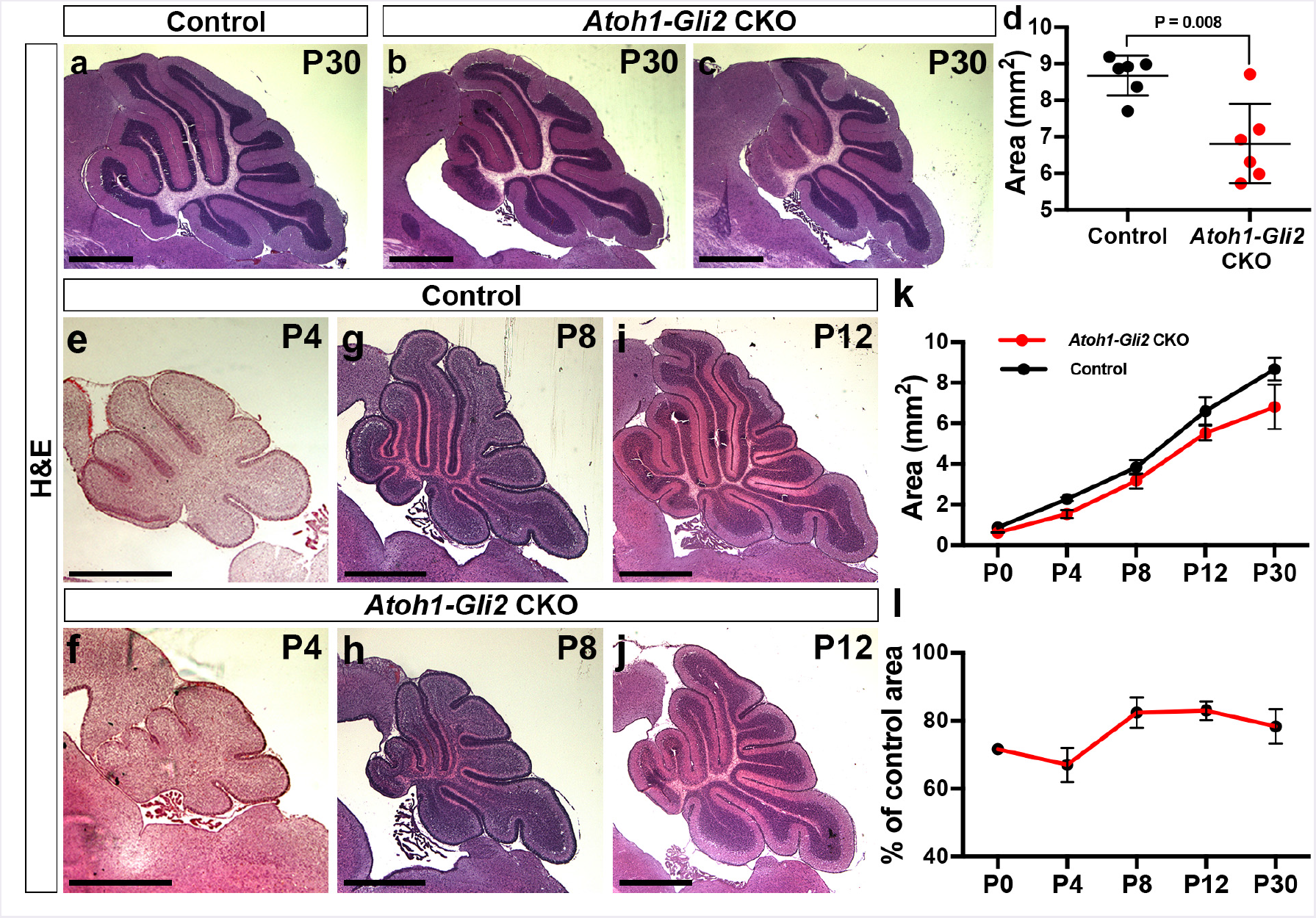
The size of the cerebellum partially recovers in *Atoh-Gli2* CKOs over time. (**a-c**) Mid-sagittal sections of P30 *Gli2^loXlox^* (control, **a**) and *Atoh1-Cre*/+; *Gli2^lox/lox^* (*Atoh1-Gli2* CKO, **b-c**) cerebellum stained with Hematoxilin and Eosin (H&E). (**d**) Graph of the area of mid-sagital CB sections of P30 *Gli2^lox/lox^* (control, black) (n=6) and *Atoh1-Cre*/+; *Gli2^lox/lox^* (*Atoh1-Gli2* CKO, red) (n=6) mice. (**e-j**) Mid-sagittal CB sections of P4 (**e-f**), P8 (**g-h**) and P12 (**i-j**) *Gli2^lox/lox^* (control, **e, g and i**) and *Atoh1-Cre*/+; *Gli2^lox/lox^* (*Atoh1-Gli2* CKO, **f, h and j**) mice stained with H&E. (**k**) Graph of the area of 3 mid-sagital sections of *Gli2^lox/lox^* (control, P0: n=3, P4: n=3, P8: n=3, P12: n=3 and P30: n=6) and *Atoh1-Cre*/+; *Gli2^lox/lox^* (*Atoh1-Gli2* CKO, P0: n=3, P4: n=2, P8: n=6, P12: n=6 and P30: n=7) cerebella. (**l**) Graph showing the decrease in area of 3 mid-sagital sections as a percentage of *Atoh1-Gli2* CKO cerebella during development. All graphical data are presented as means ± SEM and significance determined using two-tailed T-test. Scale bars represent 1mm.

In summary, we found that depletion of the EGL at P0 by removing *Gli2* from embryonic GCPs is not sufficient to induce consistent hypoplasia of the vermis at P30. This raised the possibility of a compensation mechanism that ensures the global recovery of the developing CB after genetic injury.

### Wild type cells replenish the anterior EGL of *Atoh1-Gli2* CKOs

Since the final size of the CB is largely dependent on the expansion of the EGL, we analyzed *Atoh1-Gli2* CKO cerebella at P8 when the EGL is normally thick. Similar to our previous study using irradiation at P1 (25), the EGL was replenished with proliferating cells by P8 in *Atoh1-Gli2* CKO animals. We therefore performed *in situ* hybridization (ISH) and analyzed the expression of *Gli1.* Although *Gli1* expression was greatly diminished at P0 (Fig. 1h), the EGL of P8 *Atoh1-Gli2* CKOs exhibited *Gli1* expression throughout the anterior EGL, comparable to that observed in control animals and the posterior EGL of mutants (Fig. 3a-b). In addition and unlike at P0, no difference in *Gli1* expression was observed in the PCL at P8 (Fig. 3a-b). Moreover, *Gli1+* cells expanded in the EGL, as revealed by the proliferation maker Ki67, and increased the size of the EGL by seemingly delaying their differentiation compared to controls since the proliferative (Ki67+) outer EGL [oEGL] was thicker and the inner EGL [iEGL] thinner compared to controls (n=8) (Fig. 2c-d). Interestingly, GCPs in the partially rescued anterior EGL expressed a low level of the stem cell marker SOX2 at P8, something that was never observed in control animals (Fig. 2e-f). Furthermore, although the GCPs in the EGL of *Atoh1-Gli2* CKOs expressed the EGL marker *Atoh1* (as shown by Atoh-GFP staining, Fig S4a-b), only a small subset of cells in the anterior EGL expressed *Cre* in the anterior CB compared to controls (*Atoh1-Cre*/+; *Gli2*^flox/+^ or *Atoh1-Gli2* het) (Fig. 3g-h). and Fig. S4a-b). Consistent with the present of wild type (WT) cells in the EGL, GLI2 protein was detected in the vast majority of cells in the EGL (inset Fig. 3h). Interestingly, TUNEL analysis showed an apparent increase of cell death in the replenished P8 EGL compared to controls (Fig. 4 c-d). One interpretation of these results is that WT cells replenish the EGL and then only a small subset the cells are able to turn on *Cre* expression, and those that do express *Cre* and delete *Gli2* undergo cell death (Fig. S4c-d). Thus, the reduction of GCPs in the EGL of *Atoh1-Gli2* CKOs at P0 stimulates a compensation process that leads to replenishment of the GCPs.

**Fig. 3.**
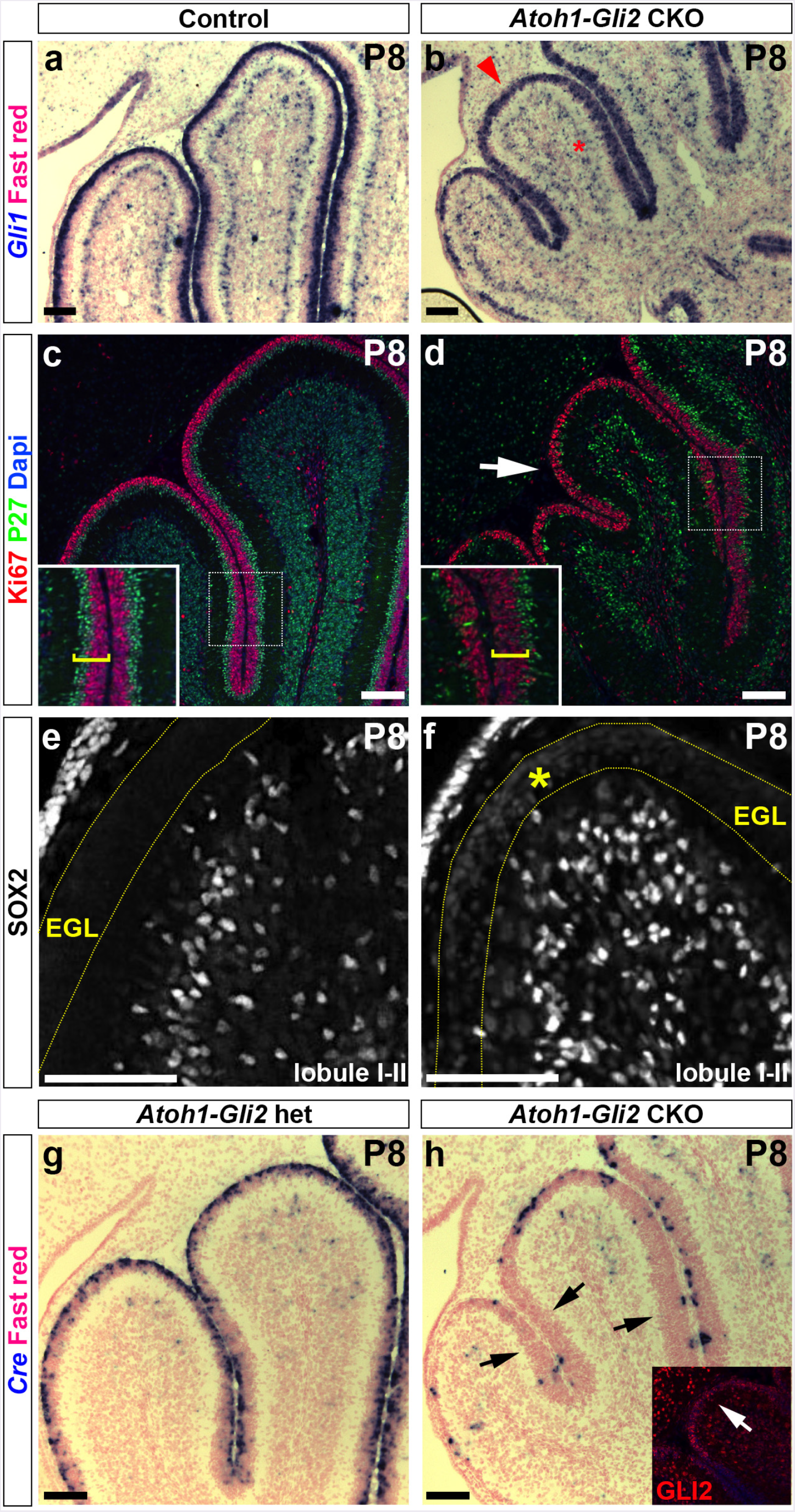
Loss of *Gli2* mutant GCPs at P0 is compensated by wild type (WT) cells at P8. (**a-b**) *In situ* RNA hybridization analysis of *Gli1* on P8 midsagittal cerebellar sections of *Gli2^lox/lox^* (control, **a**) and *Atoh1-Cre*/+; *Gli2^lox/lox^* (*Atoh1-Gli2* CKO, **b**) mice. Red arrowhead indicates the strong *Gli1* expression in the mutant EGL and red asterisks indicate normal *Gli1* expression in Bergmann glia in the Purkinje Cell Layer (PCL). (**c-f**) FIHC detection of the indicated proteins and dapi on P8 mid-sagittal cerebellar sections of *Gli2^lox/lox^* (control, **c and e**) and *Atoh1-Cre*/+; *Gli2^lox/lox^* (*Atoh1-Gli2* CKO, (**d and f**) mice. High power images are shown of the areas indicated by white rectangles in **c** and **d** with the thickness of the EGL indicated by yellow brackets. The white arrow in **d** indicates the proliferating EGL. (**e and f**) high magnification in lobule I-II region. EGL is indicated by the yellow dotted line and yellow asterisk indicates low level of SOX2 expression in the mutant EGL. (**g-h**)) *In situ* hybridization of *Cre* RNA on P8 midsagittal cerebellar sections of *Gli2^lox/lox^* (control, **g**) and *Atoh1-Cre*/+; *Gli2^lox/lox^* (*Atoh1-Gli2* CKO, **h**) mice. Black arrows indicate the loss of *Cre* expression in the partially rescued EGL. White arrow indicates the presence of GLI2 protein in the EGL (inset, **h**). Scale bars represent 100μm.

**Fig. 4.**
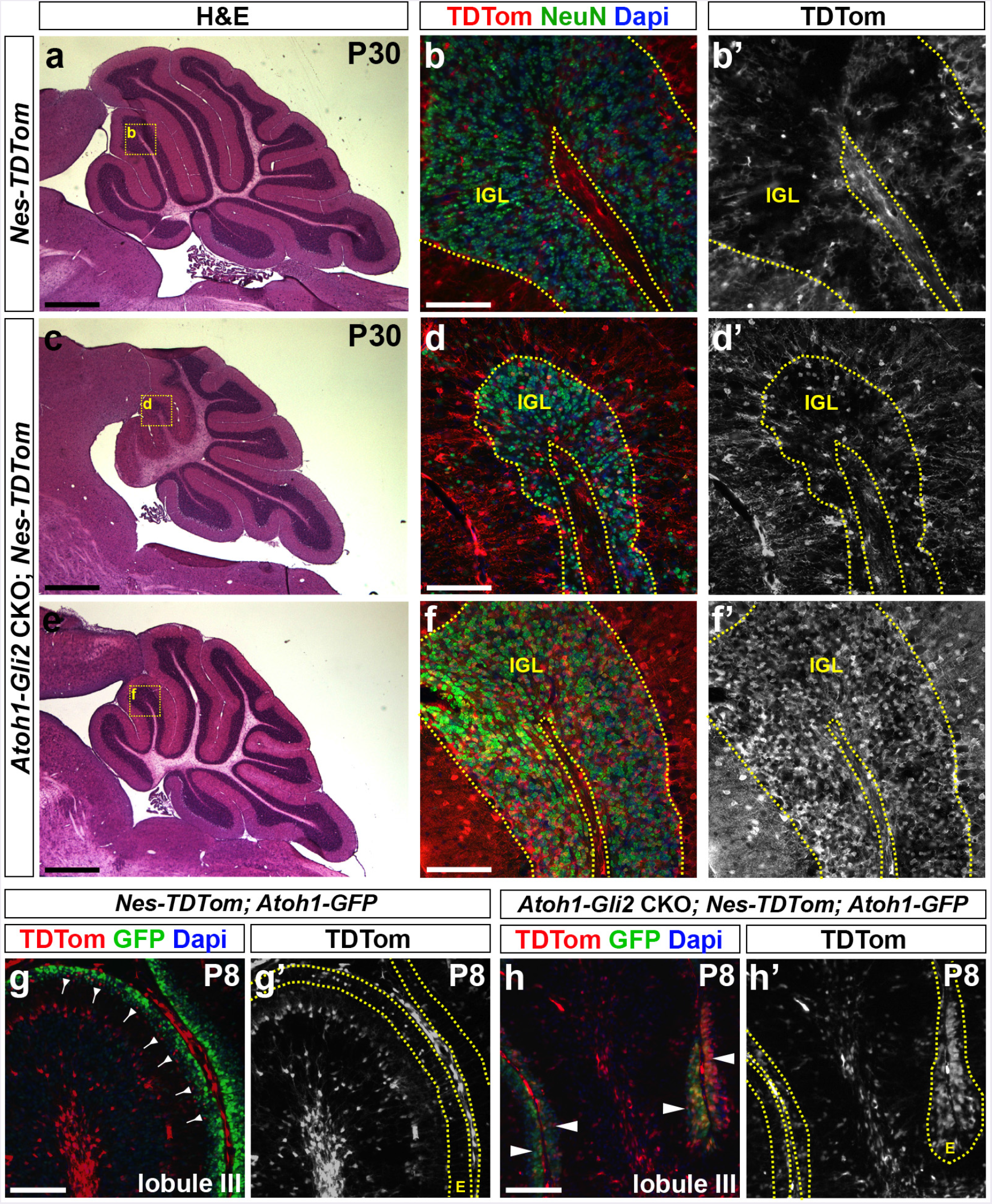
Nestin-Expressing Progenitors (NEPs) populate the EGL, express GCP markers and produce granule cells in response to loss of *Gli2* in the EGL. (**a, c and e**) H&E staining of sagittal sections of the vermis of P30 *Nes-FlpoER*/+; *R26*^*FSF-TDTom*/+^ (*Nes-TDTom*, **a**) and *Atoh1-Cre*/+; *Gli2^lox/lox^*; *Nes-FlpoER*/+; *R26*^*FSFTDTom*/+^ (*Atoh1-Gli2* CKO; *Nes-TDTom*, **c and e**) mice injected with Tm at P0. (**b, d and f**) FIHC detection of the indicated proteins and dapi on mid-sagittal cerebellar sections at P30. High power images are shown of the areas indicated by yellow rectangles in **a, c and e**. IGL is outlined by the yellow dotted line. (**g-h**) FIHC detection of the indicated proteins and dapi on mid-sagittal cerebellar sections (lobule III) of P8 *Nes-FlpoER*/+; *R26*^*FSF-TDTom*/+^; *Atoh1-GFP*/+ (*Nes-TDTom; Atoh1-GFP*, **g**) and *Atoh1-Cre*/+; *Gli2^lox/lox^*; *Nes-FlpoER*/+; *R26*^*FSF-TDTom*/+/+^; *Atoh1-GFP*/+ (*Atoh1-Gli2* CKO; *Nes-TDTom*; *Atoh1-GFP*, **h**) mice injected with Tm at P0. The EGL (E) is outlined by the yellow dotted lines. Backward arrows indicate TDTom+ and Atoh1-GFP-cells in the inner EGL. White arrowheads indicate TDTom+ and Atoh1-GFP+ cells in the inner EGL. Scale bars represent 1mm (**a, c and e**) and 100μm (**b, d, f, g and h**).

### NEPs switch their fate to become GCPs and produce GCs in *Atoh1-Gli2* CKO cerebella

Our previous study demonstrated that SOX2+ NEPs can replenish the EGL after injury (25), and our present results revealed that the rescued EGL in P8 *Atoh1-Gli2* CKO cerebella expresses a low level of SOX2. We thus hypothesized that WT NEPs are able to change their fate to become GCPs and replenish the EGL as part of a compensation mechanism. Using a *Nestin-FlpoER* transgene (40) and a Flippase (Flp)-dependent *R26* reporter allele that expresses TDTom, we performed genetic inducible fate mapping (GIFM) of NEPs in the *Atoh1-Gli2* CKO cerebella. In contrast to P30 controls (*Nestin-FlpoER*/+; *R26*^*frt-STOP-frt-TDTom*/+^ or *Nes-TDTom* mice administrated Tm at P0), *Atoh1-Gli2* CKO mutants (n=6) had an extensive contribution of TDTom+ cells to the NeuN+ GC population in the IGL (Fig. 4a-f). Interestingly, the degree of recovery of the vermis in P30 *Atoh1-Gli2* CKO cerebella correlated with the contribution of NEP-derived TDTom+ cells in the IGL (compare (Fig. 4c-d). to 4e-f). Similar results were obtained when analyzing the hemispheres (Fig. S5). However, and consistent with our analysis of CB size, there appeared to be less variability in the percentage of TDTom+ cells observed in the hemispheres compared to the vermis. Analysis of the vermis at P8 showed an increase in the number of Nestin-derived TdTom+ cells in the EGL compared to controls (Fig. 4g-h). Furthermore, TDTom+ cells in the P8 EGL already expressed the GCP marker *Atoh1* (as shown by Atoh1-GFP staining) (Fig 4g-h).

Taken together, our findings indicate that in *Atoh1-Gli2* CKOs in which the EGL is depleted at P0 NEPs repopulate the EGL, turn on EGL genes (*Atoh1*) and down-regulate NEP genes (SOX2) and then differentiate into GCs that populate the IGL.

### A subset of proliferating PCL NEP-derived cells migrate to the IGL in *Atoh1-Gli2* CKO cerebella

We next analyzed the behavior of NEPs using a *Nes-CFP* reporter line (43). Consistent with our GIFM experiment and unlike control animals *(Nes-CFP)*, the EGL of *Atoh1-Gli2* CKOs expressed a high level of CFP (Fig. 5a-b). Surprisingly, streams of CFP+ cells were seen in lobule 3 spanning between the IGL and EGL that were not present in controls (Fig. 5a-b).) or in irradiated mice (25). Interestingly, some cells in the streams expressed the proliferation maker Ki67 as well as the GCP/GC marker PAX6 (Fig. 5c-d).) suggesting that a subset of NEP-derived cells were not able to stay in the EGL and thus migrated back to the cerebellar cortex. To test this idea we performed live imaging of P8 *Nes-CFP* cerebellar slice cultures from both control and *Atoh1-Gli2* CKOs (*Atoh-Gli2* CKO; *Nes-CFP*). Strikingly, by tracking the movement of individual cells during ~6hrs of imaging we observed Nes-CFP+ cells actively migrating from the PCL to the EGL in slices from *Atoh-Gli2* CKO cerebella but not control mice at P8 (Fig. 5e-f). and sup. videos 1-2). The CFP+ layer of cells also appeared thicker in the mutants, indicating the NEPs expanded in number. Interestingly, in the areas containing streams of CFP+ cells the majority of cells that were tracked moved in the opposite direction from the EGL to the IGL (Fig. 5g and sup. video 3). Our live imaging experiments thus provide evidence that NEPs located in the PCL expand and then migrate to replenish the EGL in response to GCP-specific loss of *Gli2*. Furthermore, a subset of NEP-derived cells is not able to integrate into the EGL and migrate back down to towards the cerebellar cortex.

**Fig. 5.**
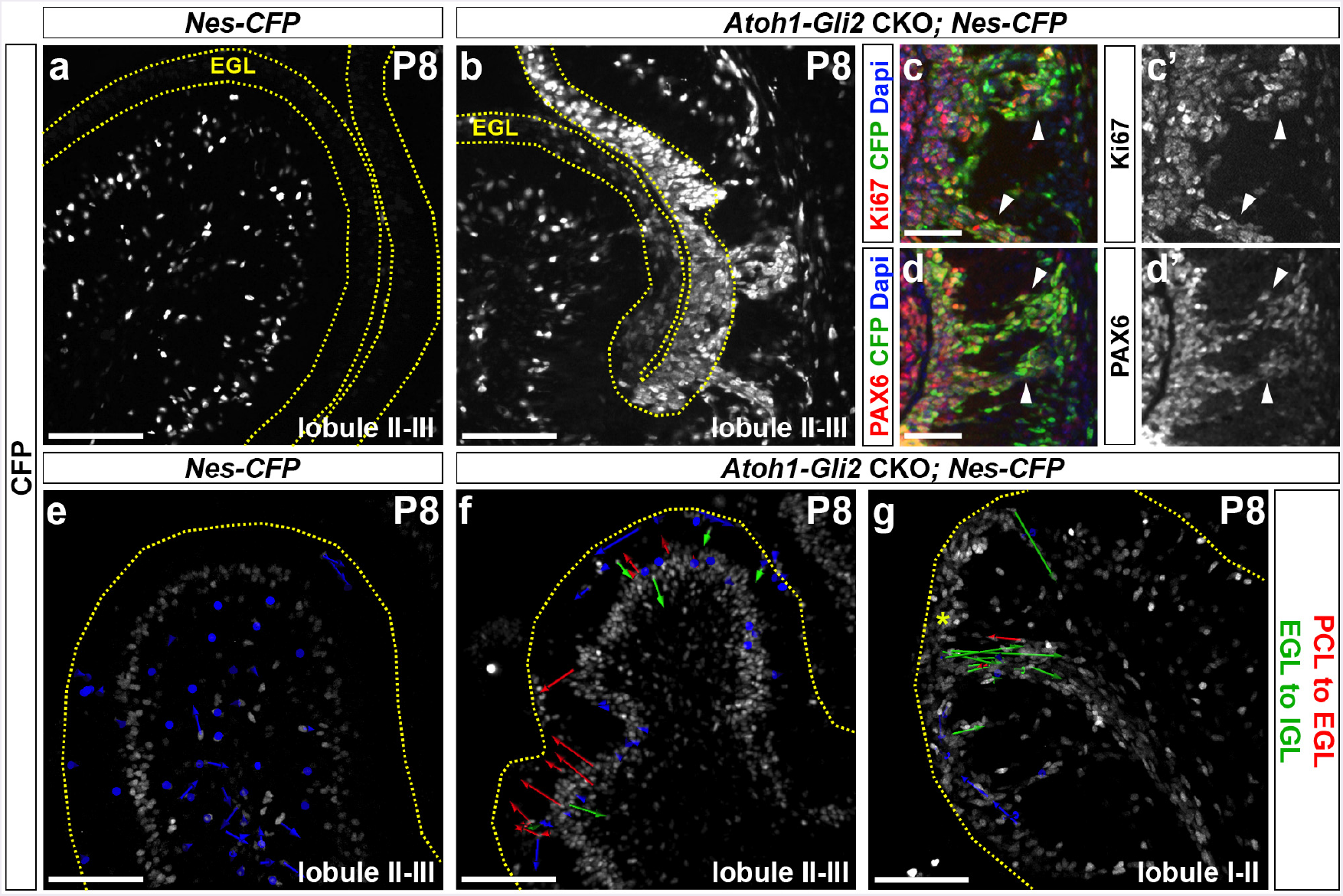
PCL NEPs migrate to the EGL and a subset of proliferating NEP-derived GCP-like cells migrate back into the cerebellar cortex in streams. (**a-b**) FIHC detection of CFP protein on mid-sagittal cerebellar sections (lobule II-III) of P8 *Nes-CFP*/+ (*Nes-CFP*, **a**) and *Atoh1-Cre*/+; *Gli2^lox/lox^*; *Nes-CFP*/+ (*Atoh1-Gli2* CKO; *Nes-CFP*, **b**) mice. The EGL is indicated by the yellow dotted lines. (**c-d**) FIHC detection of the indicated proteins and dapi on high magnifications focusing on CFP+ streams of mid-sagittal cerebellar sections of P8 *Atoh1-Cre*/+; *Gli2^lox/lox^*; *Nes-CFP*/+ (*Atoh1-Gli2* CKO; *Nes-CFP*) mice. White arrowheads indicate co-localization of CFP with the indicated protein. (*e-g*) Detection of native CFP fluorescence on sagittal slice cultures of the vermis (lobule II-III, **e and f** and lobule I-II, **g**) of P8 *Nes-CFP*/+ (*Nes-CFP*, **e**) and *Atoh1-Cre*/+; *Gli2^lox/lox^*; *Nes-CFP*/+ (*Atoh1-Gli2* CKO; *Nes-CFP*, **f and g**) mice showing displacement of CFP+ cells during 6h of imaging. Arrow color code is as indicated. The upper edge of the EGL is indicated by a yellow dotted line. Scale bars represent 100μm (**a-b and e-g**) and 50μm (**c-d**).

### *Gli2* CKO in NEPs inhibits the recovery of the EGL in *Atoh1-Gli2* CKOs

Since we have shown previously that SHH signaling (*Smo*) is necessary in NEPs for CB recovery following irradiation, we tested whether *Gli2* plays a role in this process. We generated littermates of 4 different genotypes that were administered Tm at P0, and analyzed each genotype (n=3) at P8, P12 and P21: *Gli2^flox/flox^* WT (control) mice, *Nestin-FlpoER*/+; *R26*^*MASTR*^+; *Gli2*^*flox/flox*^ single (*Nes-Gli2* CKOs mutants lacking *Gli2* in NEPs), *Atoh1-Cre*/+; *Gli2*^*floxflox*^ single (*Atoh1-Gli2* CKOs lacking *Gli2* in anterior GCPs) and *Nestin-FlpoER*/+; *R26*^*MASTR*/^+; *Atoh1-Cre*/+; *Gli2*^flox/flox^ double (*Atoh1-Nes-Gli2* CKOs lacking *Gli2* in NEPs and GCPs) mutants. We did not observe any obvious phenotype in the *Nes-Gli2* CKOs at all stages compared to controls (compare Fig. 6c-d). to a-b, Fig. S6c-d to a-b and Fig. S7d to a). However, H&E staining revealed that the anterior CB was greatly reduced in the double mutants (*Atoh1-Nes-Gli2* CKOs) compared to *Atoh1-Gli2* CKOs at both P8 and P12 (compare Fig. 6e to g and Fig. S6e to g). Analysis of proliferation (Ki67) and differentiation (P27) markers at both stages showed not only that the anterior EGL was greatly depleted in *Atoh1-Nes-Gli2* CKO compared to *Atoh1-Gli2* CKO cerebella, but also that *Atoh1-Nes-Gli2* CKO animals failed to form a proper P27+ inner EGL and IGL (compare Fig. 6f to h and Fig. S6f to h). Analysis of the fate of GFP+ Nestin-expressing cells in P21 *Atoh1-Nes-Gli2* CKOs using a Flp-mediated *R26* reporter allele that expresses βGalactosidase (Bgal)(*R26*^*frt-STOP-frtlacZ/+*^) revealed that unlike *Atoh1-Gli2* CKOs in which Nestin-derived GCs populated the IGL ((Fig. 4a-f). and Fig. S7c), the *Gli2* mutant Nestin-derived cells did not populate the IGL (Fig. S7f). The few GFP+ cells in the IGL of the double mutants were likely interneurons or astrocytes. These results demonstrate that SHH-signaling through GLI2 plays a crucial role during NEP-mediated cerebellar recovery from loss of GCPs.

**Fig. 6.**
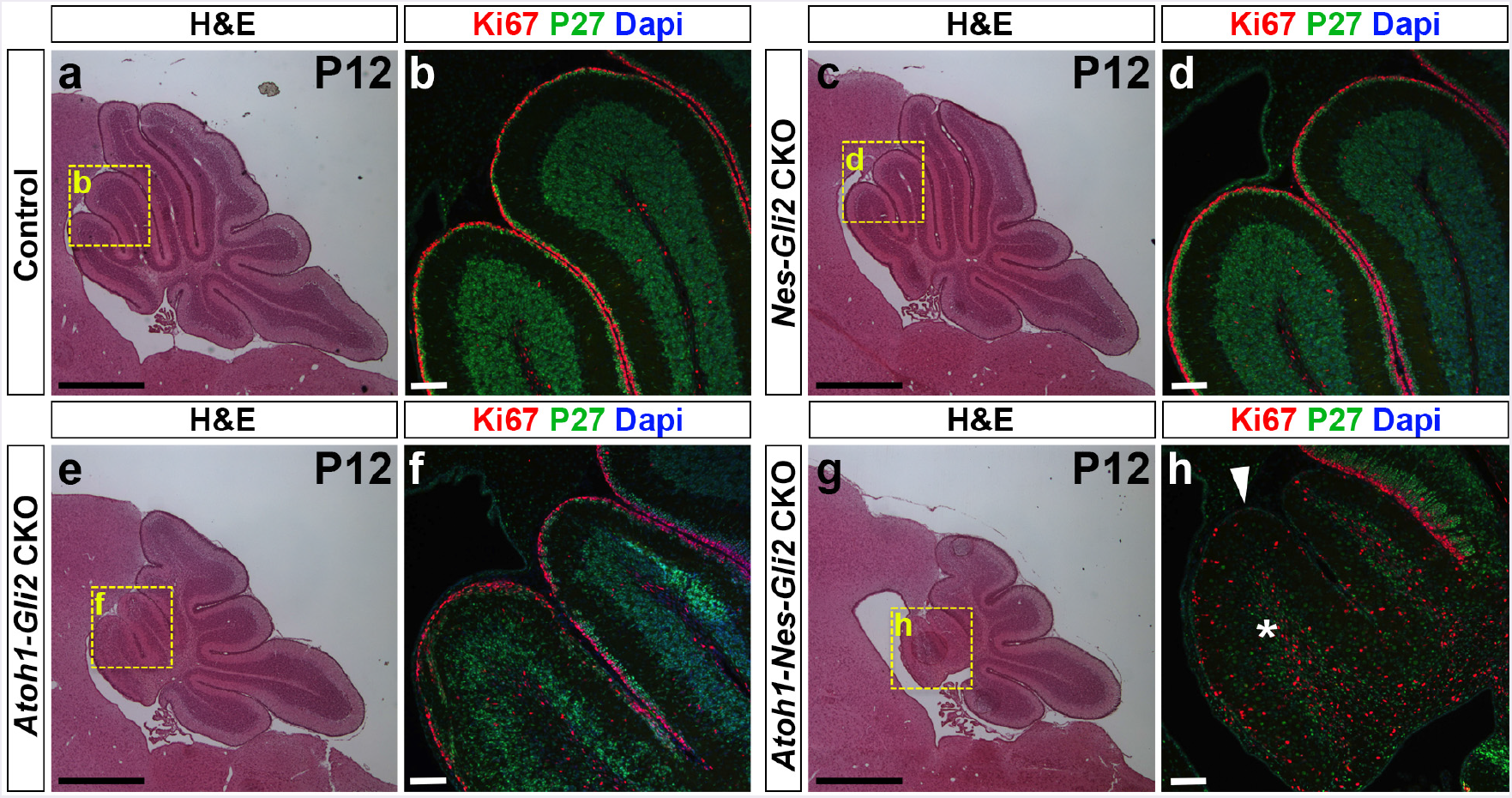
Inactivation of *Gli2* in both *Nestin-* and Atoh7-expressing cells inhibits the recovery of the CB compared to in mice lacking *Gli2* only in the EGL. (**a, c, e and g**) H&E of sagittal sections of the cerebellar vermis of P12 *Gli2^flox/flox^* (Control, **a**), *Nes-FlpoER*/+; *R26*^*MASTR*/+^; *Gli^flox/flox^* (*Nes-Gli2* CKO, **c**), *Atoh1-Cre*/+; *Gli^flox/flox^* (*Atoh1-Gli2* CKO, **e**), and *Atoh1-Cre*/+; *Nes-FlpoER*/+; *R26*^MASTR/+^; *Gli^flox/flox^* (*Atoh1-Nes-Gli2* CKO, **g**) mice injected with Tm at P0. Note that inactivation of *Gli2* only in *Nestin-* expressing cells has no major effect of CB development at P12. However, inactivation of *Gli2* in *Nestin*-expressing cells inhibits the compensation mechanism when *Gli2* is removed in GCPs (**g** compared to **e**). (**b, d, f and h**) High magnifications (as shown by yellow squares in **a, c, e and g**) of anterior vermis of P12 *Gli2^fox/fox^* (**b**), *Nes-FlpoER*/+; *R26*^*MASTR*/+^; *Gli^flox/flox^* (*Nes-Gli2* CKO, **d**), *Atoh1-Cre*/+; *Glif^ox/flox^* (*Atoh1-Gli2* CKO, **f**), and *Atoh1-Cre*/+; *Nes-FlpoER*/+; *R26*^*MASTR*/+^; *Gli^flox/flox^* (*Atoh1-Nes-Gli2* CKO, **h**) cerebella stained with the indicated proteins and dapi. White arrowhead and white asterisk indicate the loss of EGL and IGL respectively in *Atoh1-Nes-Gli2* CKO. Scale bars represent 1mm (**a, c, e and g**) and 100μm (**b, d, f and h**).

## Discussion

In this study we developed a conditional mutant strategy to delete *Gli2* (the gene encoding the major effector of SHH signaling) in the vast majority of GCPs using a *Atoh1-Cre* transgene that is first expressed in embryonic GCPs (45). Although we show that SHH-GLI2 signaling is crucial for generating the correct pool of GCs, deletion of *Gli2* in the EGL using this transgene is not sufficient to induce a major hypoplasia of the adult CB. We discovered that although as expected the GCP pool was greatly diminished at birth, it was subsequently replenished due to adaptive reprogramming of WT NEPs, and since the transgene did not turn on in many of the newly generated GCPs, the EGL recovered and generated GCs.

SHH regulates a variety of cell behaviors depending on the cellular context and concentration of SHH ligand (29). We demonstrated that SHH signaling through the GLI2 activator not only influences proliferation of GCPs by keeping them in an undifferentiated state and increasing their proliferation rate, but also maintains their survival. Our previous study showed that massive killing of GCPs at an early stage of postnatal development (P2-3) triggers NEP-dependent recovery of the EGL (25). It is therefore possible that cell death is involved in triggering NEPs to change their fate and generate GCPs, possibly because an alarm signal is released. An alternative mechanism for the stimulation of NEPs is that PCs are able to sense the EGL injury because of a lack of excitatory inputs from parallel fibers of GCs, and as part of the response PCs modulate the amount of HH signaling in the PCL NEPS by concentrating the ligand at their surface (25). Cell death in the EGL *per se* would therefore not be the main trigger that recruits NEPs to the EGL but instead the lack of differentiation of GCPs would stimulate NEPs. Consistent with a role for GCs in regulating NEP behaviors, the *Atoh1-Cre* transgene is first expressed in GCPs at E13.5 (17), but the replenishment of the EGL in *Atoh1-Gli2* CKOs only occurs several days after the EGL is depleted and when the layer of differentiating GCs in the EGL and IGL of control animals first becomes apparent (P3-P4). Thus, the possible involvement of NEPs in compensation processes should be consider when analyzing the phenotype of conditional mutants that affect not only trophic factors that stimulate GCP proliferation and survival, but also genes involved in differentiation and migration of GCPs.

Our study clearly demonstrates that depletion of GLI2 in the EGL using an *Atoh1-Cre* transgene is not sufficient to drastically and consistently reduce the size of the adult CB. Curiously, *Atoh1-CreER/+; Smo^loxP/Δ^* mice in which one allele of the *Smoothened* gene was deleted in the germline and the deletion was dependent on tamoxifen (Tm) inducible Cre *(Atoh1-CreER)* showed a severe CB hypoplasia (13). A possible explanation for the different phenotype from *Atoh1-Gli2* CKOs lies in our finding that HH signaling is crucial for the expansion of PCL NEPs and their migration to the EGL for effective recovery (25). In addition, we found that Tm diminishes CB recovery by delaying the response of NEPs after injury. Thus the combination of a lower level of SMO protein in NEPs as well as administration of Tm to *Atoh1-CreER/+; Smo^loxP/Δ^* mice might have blunted the response of NEPs to Smo-dependent depletion of the EGL, resulting in little compensation and a large decrease in size of the CB compared to controls.

We have uncovered that adaptive reprogramming of NEPs occurs in *Atoh1-Gli2* CKO animals. However, we observed a large variability in the adult CB phenotype, especially in the vermis, suggesting that some cerebella are not able to efficiently recover after *Gli2* depletion in the EGL. Although ATOH1+ WT cells replenish the EGL over time in *Atoh1-Gli2* CKOs, we found that only a variable subset of GCPs turned on the *Atoh1-Cre* transgene in different mice. Consistent with *Atoh1-Cre* deleting *Gli2* in a subset of NEP-derived GCPs, we observed an increase in TUNEL staining in the EGL of P8 *Atoh1-Gli2* CKO cerebella. It is likely that the increase in cell death negatively impacts on the expansion of new GCPs in the EGL. In addition, our live imaging experiment revealed that proliferative Nes-CFP+ and PAX6+ cells migrated from the EGL to the cerebellar cortex. We hypothesize that a factor is missing in NEP-derived GCPs that lose GLI2 after entering the EGL that is needed to maintain them in the EGL. An interesting candidate signaling pathway is SDF1-CXCR4 signaling, since *Sdf1 (Cxcl12)*, the gene encoding a well-known chemo-attractant, is specifically expressed in the meninges covering the CB (46) and two receptors, CXCR4 and CXCR7, are expressed in GCPs and the PCL, respectively (47, 48). Furthermore, SDF1 is required to maintain GCPs in the outer EGL and to make them more permissive to SHH-dependent proliferation by inhibiting cAMP or PKA activity (46). Furthermore, SHH signaling induces the transcription of *Cxcr4* and *Cxcr7*, which contain GLI binding sites in their promoters (49). We propose that in *Atoh1-Gli2* CKOs a subset of PCL NEPs are able to express CRE protein after integrating in the EGL, and the subsequent deletion of GLI2 protein reduces the expression of CXCR4, leading to migration of proliferating NEP-derive GCP-like cells back into the cerebellar cortex. Together, the increase of cell death in the EGL and the premature migration of proliferating NEP-derived GCPs could explain the variable recovery of the CB in *Atoh1-Gli2* CKOs.

The cerebellum is broadly divided along the medio-lateral axis into a central vermis and two lateral hemispheres (19). Although recovery from depletion of the EGL at P0 occurred in both regions, the compensation was more variable and less pronounced in the vermis compared to the hemispheres, with no statistical difference in hemispheric size compared to controls. Moreover the hemispheres of *Atoh1-Gli2* CKOs exhibited extra folia at P8 (arrow in FigS3), highlighting a differential recovery response along the medio-lateral axis. The vermis and hemispheres are molecularly distinct on the basis of gene expression patterns and functionally distinct based on afferent circuits (19, 50). Moreover we recently reported that hemispheric GCPs have a higher sensitivity to high level SHH-signaling than those in the vermis, and this likely contributes to the high incidence of medulloblastoma tumors in the hemispheres (50). Furthermore, since SHH signaling is required for recovery of the EGL after injury by stimulating the expansion of PCL NEPs and their migration towards the EGL (25), we propose that hemispheric NEP-derived GCPs in the EGL maintain a higher level of SHH signaling and therefore expand more rapidly and efficiently than those in the vermis.

## Conclusion

In this study we uncovered that the plasticity of cerebellar NEPs and their ability to repopulate GCPs after post-natal cerebellar loss of the EGL can occur in conditional mouse mutants. A mild phenotype in EGL-specific conditional mutants therefore does not necessarily mean a gene does not play a major role in development of the GC-lineage. Thus, the possible contribution of adaptive reprogramming of WT NEPs in a recovery process must be consider when analyzing and interpreting cerebellar phenotypes. This is particularly the case if the *Atoh1-Cre* transgene, which is broadly used in the CB field, is used to generate conditional mutants. Our findings also raise the question of whether similar recovery phenomena occur in other regions of the brain, and depending on the transgene used could complicate interpretation of mutant phenotypes.

## Acknowledgments

We thank the past and present members of the laboratory for helpful discussions during the course of our study.

## Funding

This work was supported by grants from the Brain Tumor Center at MSKCC and from the Philippe Foundation (to A.W), and from the NIH (R37-MH085726 and R01 NS092096 to A.L.J and F32 NS086163 to A.K.L.) and a National Cancer Institute Cancer Center Support Grant (P30 CA008748-48 to A.L.J).

## Availability of data and materials

All mouse lines are available from the Joyner lab or Jackson laboratories. All data generated or analyzed for this study are included in this published article.

## Authors’ contribution

A.W. and A.L.J. conceived the project; A.W., A.K.L and A.L.J. designed the research; A.W., M.M., A.K.L and D.N.S. performed the experiments; A.W., M.M., A.K.L and A.L.J. analyzed the data and all authors discussed the data; A.W. and A.L.J. wrote the manuscript with contributions from all authors.

## Competing interests

The authors declare no conflict of interest.

## Consent for publication

All authors read and approved the manuscript.

## Ethics approval and consent to participate

All animal procedures were performed according to a protocol (07-01-001) approved by the Memorial Sloan Kettering Cancer Center’s Institutional Animal Care and Use Committee.

## Supplementary material

**Fig. S1.**
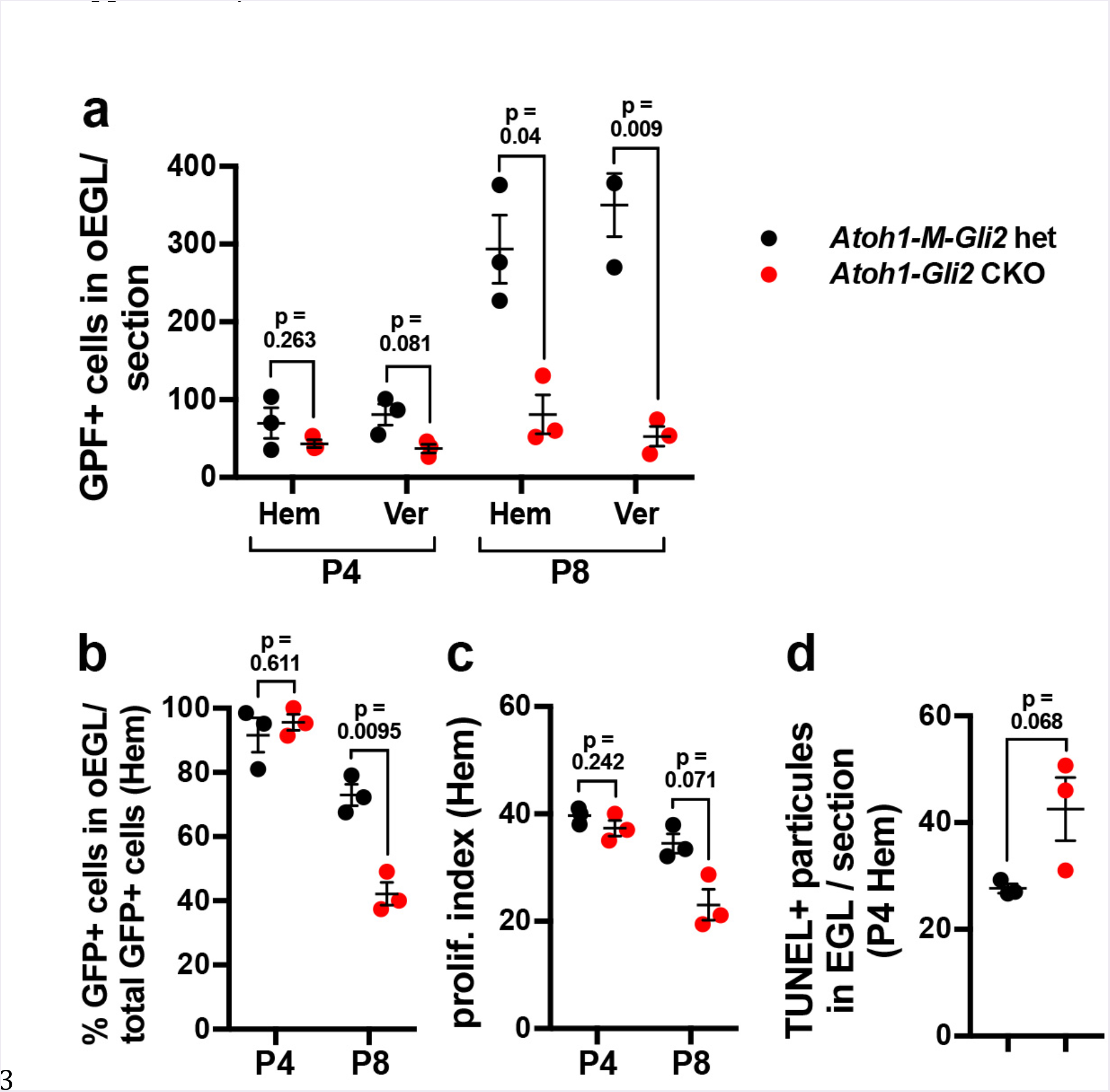
Similar to vermis, SHH/Gli2 maintains GCP in an undifferentiated state and promotes their survival in the hemisphere. (**a**) Graphs of the number of GFP+ cells in in outer (o) at P4 (n=3) and P8 (n=3) in both hemisphere and vermis of *R26*^*MASTR*/+^; *Atoh1-FlpoER*/+; *Gli2*^*lox*/+^ (*Atoh1-M-Gli2* het, black) and *R26*^MASTR^+; *Atoh1-FlpoER*/+; *Gli2^lox/lox^* (*Atoh1-M-Gli2* CKO, red) mice treated Tm at P2 (**b-d**) Graphs of the proportion of CFP+ cells in in outer (**o**) EGL at P8 (n=3) (**b**), the proliferation index at P8 (% [GFP+ EdU+] cells of all [GFP+] cells in the oEGL) (n=3) (**c**) and the number TUNEL+ particles per section at P4 (n=3) (**d**) in the hemisphere of *R26*^*MASTR*/+^; *Atoh1-FlpoER*/+; *Gli2*^*lox*/+^ (*Atoh1-M-Gli2* het, black) and *R26*^*MASTR*/^; *Atoh1-FlpoER*/+; *Gli2*^*lox/lox*^ (*Atoh1-M-Gli2* CKO, red) mice treated Tm at P2. All of the analyses were performed on 3 sections per region and per brain. All graphical data are presented as means ± SEM and significance determined using two-tailed.

**Fig. S2.**
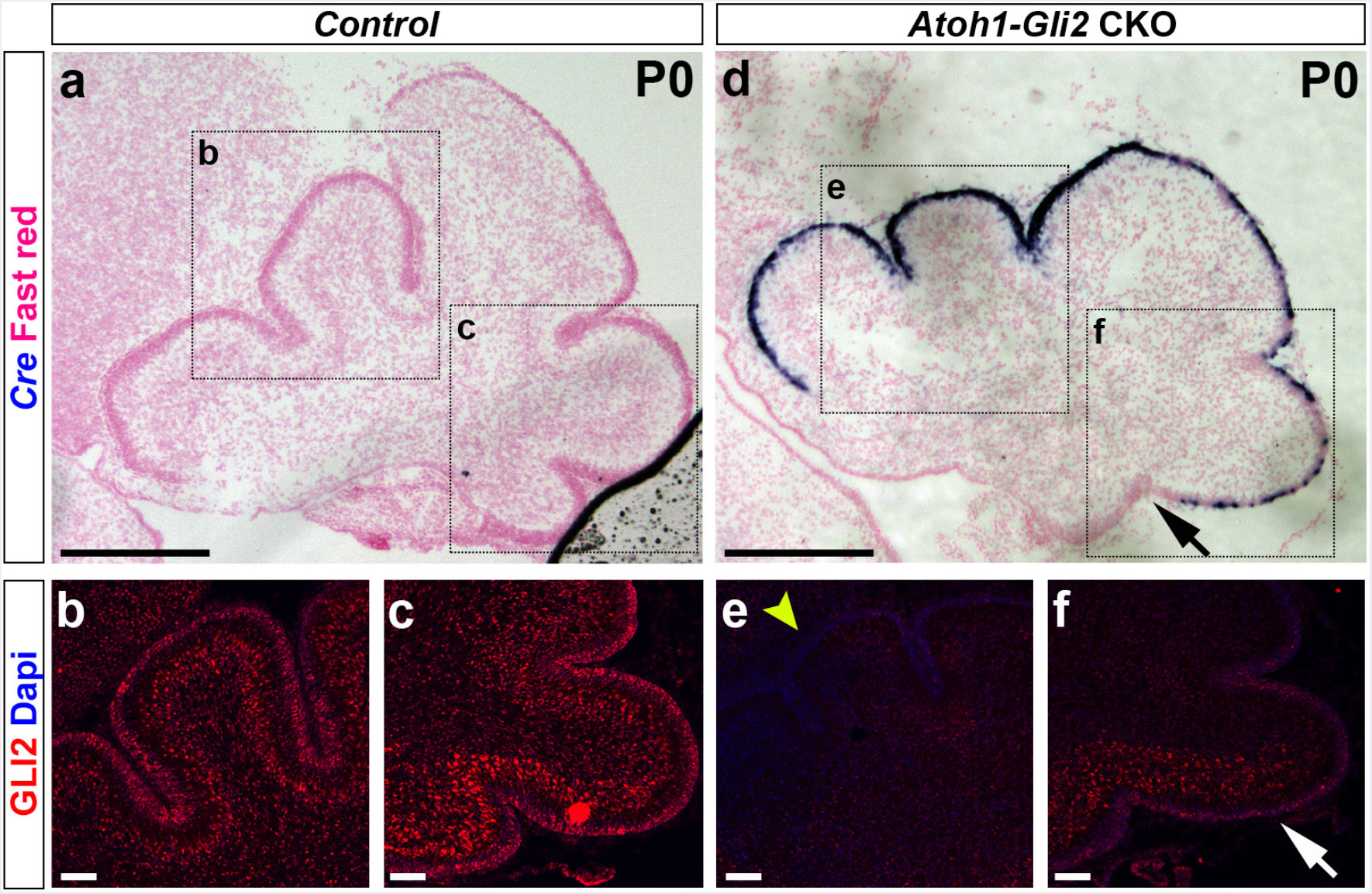
GLI2 protein is lost in P0 *Atoh1-Gli2* CKO EGL. (**a and d**) *In situ* hybridization of *Cre* mRNA on P0 mid-sagittal cerebellar sections of *Gli2^lox/lox^* (control, **a**) and *Atoh1-Cre*/+; *Gli2^lox/lox^* (*Atoh1-Gli2* CKO, **d**) mice. Black arrows indicate the lack of *Cre* expression in the most posterior part of the CB. (**b-c and e-f**) FIHC detection of GLI2 protein and dapi in the indicated regions (as shown by black squares in **a and d**) in P0 *Gli2^lox/lox^* (control, **b-c**) and *Atoh1-Cre*/+; **Gli2^lox/lox^** (*Atoh1-Gli2* CKO, **e-f**) CB. Yellow arrowhead in **e** and white arrow in **F** indicate respectively the absence and presence of GLI2 protein in the EGL. Scale bars represent 1mm (**a and d**) and 100μm (**b-c and e-f**).

**Fig. S3.**
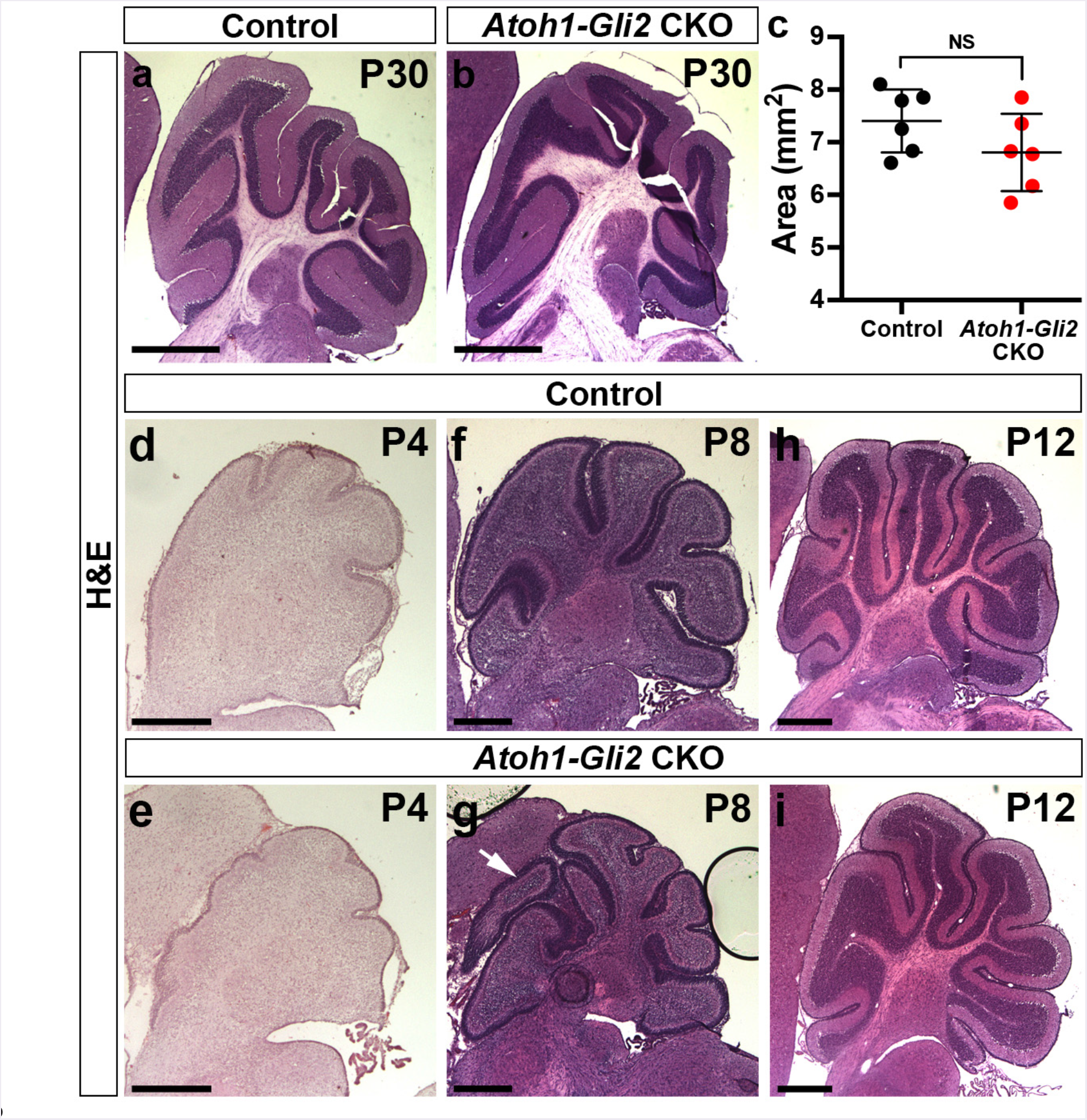
Hemisphere recovers better that vermis in *Atoh1-Gli2* CKO mice. (**a-b**)hemispheric sagittal sections of P30 *Gli2^lox/lox^* (control, **a**) and *Atoh1-Cre*/+; *Gli2^lox/lox^* (*Atoh1-Gli2* CKO, **b**) cerebellum stained with Hematoxilin/Eosin (H&E). (**C**) Graph of the area of hemispheric sagital sections of P30 *Gli2^lox/lox^* (control, black) (n=6) and *Atoh1-Cre*/+; *Gli2^lox/lox^* (*Atoh1-Gli2* CKO, red) (n=6) CB. (**d-i**) Hemispheric sagittal sections of P4 (**d-e**), P8 (**f-g**) and P12 (**h-i**) *Gli2^lox/lox^* (control, **d, f and h**) and *Atoh1-Cre*/+; *Gli2^lox/lox^* (*Atoh1-Gli2* CKO, **e, g and i**) cerebellum stained with H&E. White arrow indicates the presence of extra folia. Scale bars represent 1mm (**a-b**) and 500μm (**b-c and d-i**).

**Fig. S4.**
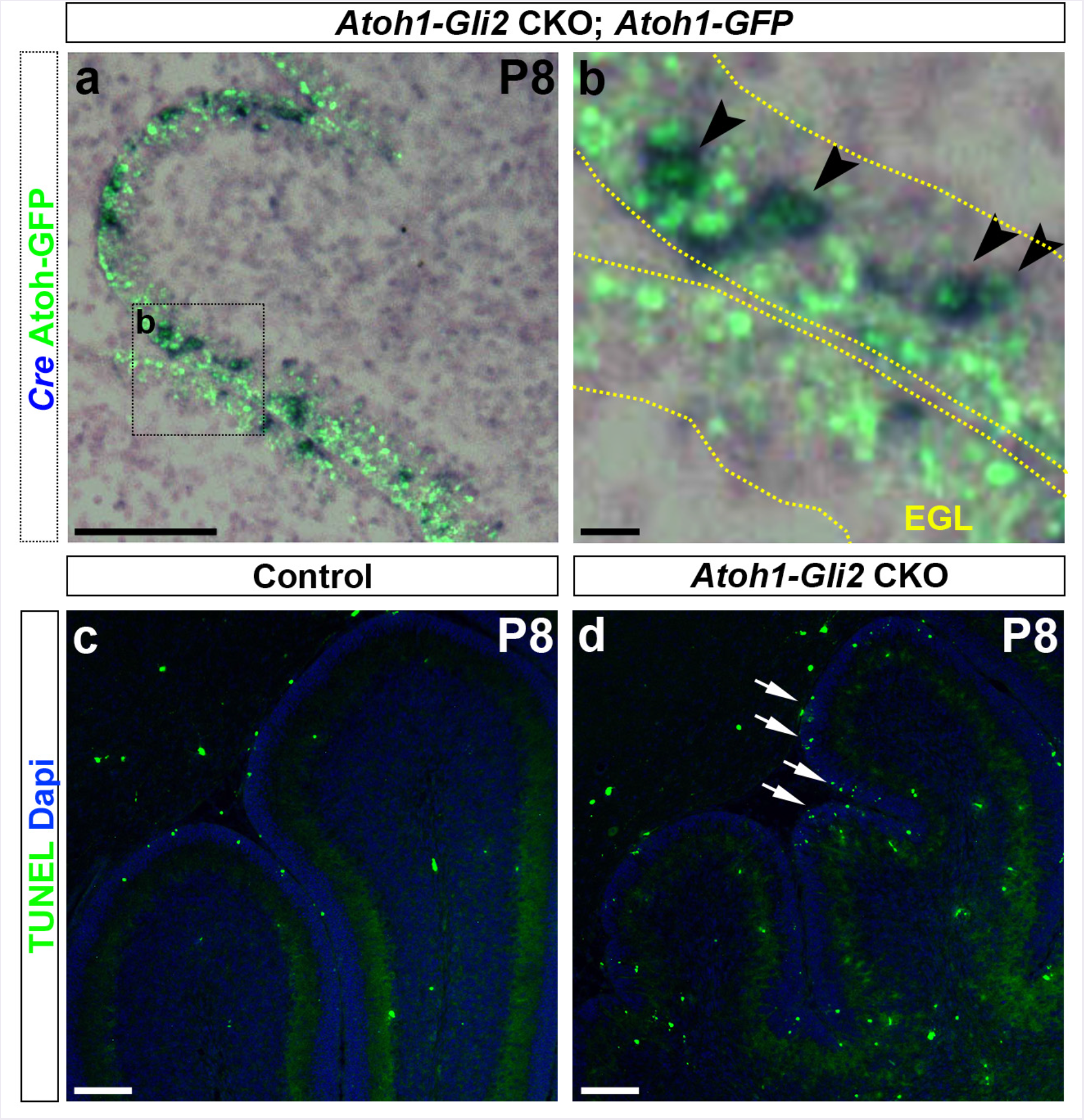
Rescued EGL still exhibits an increase in cell death. (**a-b**) Detection of native GFP fluorescence and *In situ* hybridization of *Cre* mRNA on mid-sagittal section (lobule II-III) of P8 *Atoh1-Cre*/+; *Gli2^lox/lox^*; *Atoh1-GFP*/+ (*Atoh1-Gli2* CKO; *Atoh1-GFP*) mice. High power image is shown of the area indicated by black rectangles in **a**. in **b**, EGL is indicated by the yellow doted line and black arrowheads indicate ATOH1-GFP+/ *Cre*+ cells, (**c-d**) TUNEL and dapi detection on mid-sagittal sections of P8 *Gli2^lox/lox^* (Control) and *Atoh1-Cre*/+; *Gli2^lox/lox^* (*Atoh1-Gli2* CKO) CB. White arrows indicate the presence of in the EGL (**d**). Scale bars represent 100μm (**a and c-b**) and 10μm (b).

**Fig. S5.**
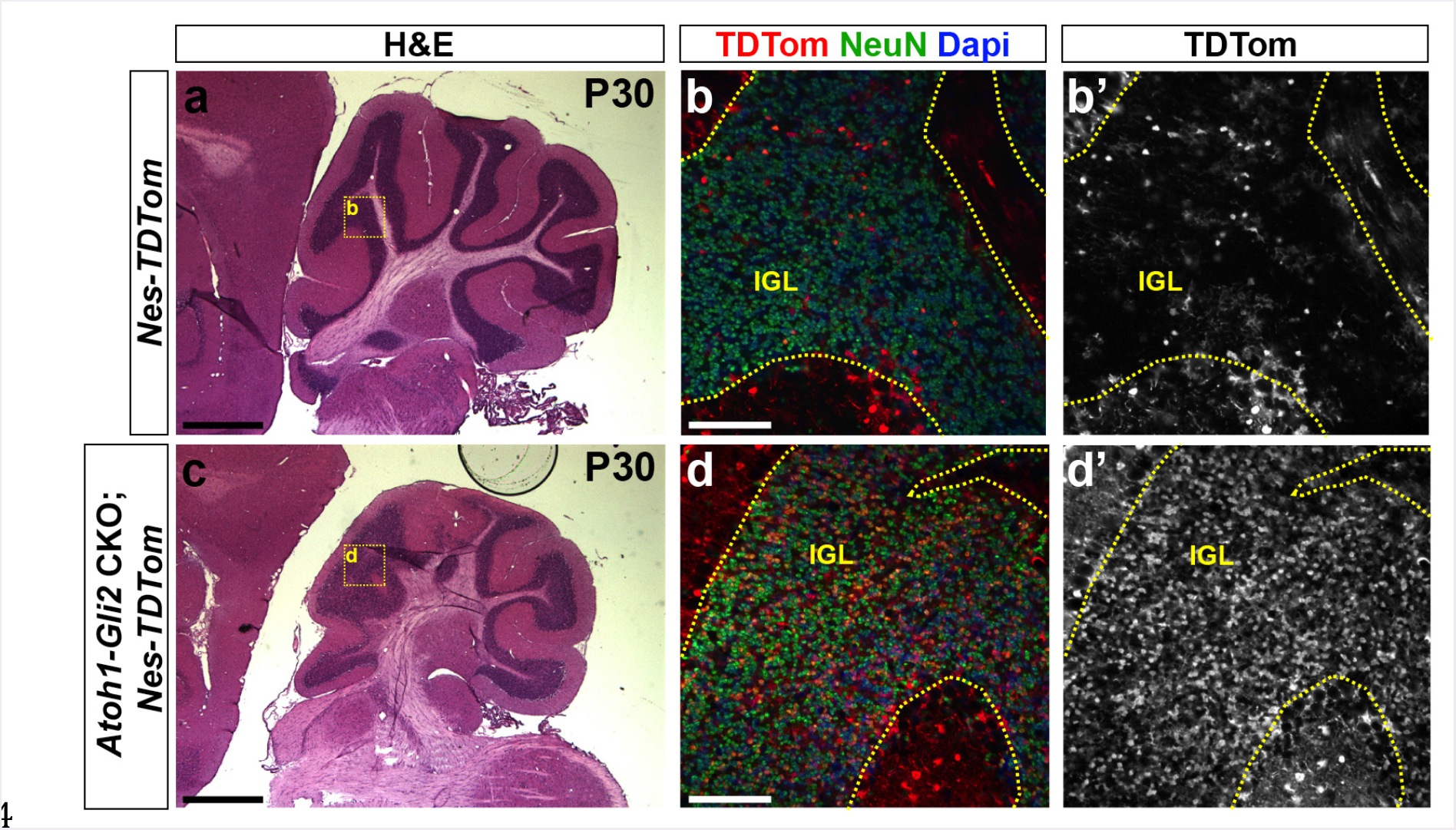
*Nestin*-Expressing Progenitors (NEPs) differentiate into Granule neurons in response to loss of *Gli2* in the hemisphere. (**a and c**) H&E of hemispheric sagittal sections of P30 *Nes-FlpoER*/+; *R26*^*FSF-TDTom*/+^ (*Nes-TDTom*, **a**) and *Atoh1-Cre*/+; *Gli2^lox/lox^*; *Nes-FlpoER*/+; *R26*^*FSF-TDTom*/+^ (*Atoh1-Gli2* CKO; *Nes-TDTom*, **c**) mice injected with Tm at P0. (**b and d**) FIHC detection of the indicated proteins and dapi on hemispheric sagittal cerebellar sections at P30. High power images are shown of the areas indicated by yellow rectangles in (**a and c**). IGL is indicated by the yellow doted line. Scale bars represent 1mm (**a and c**) and 100μm (**b and d**).

**Fig. S6.**
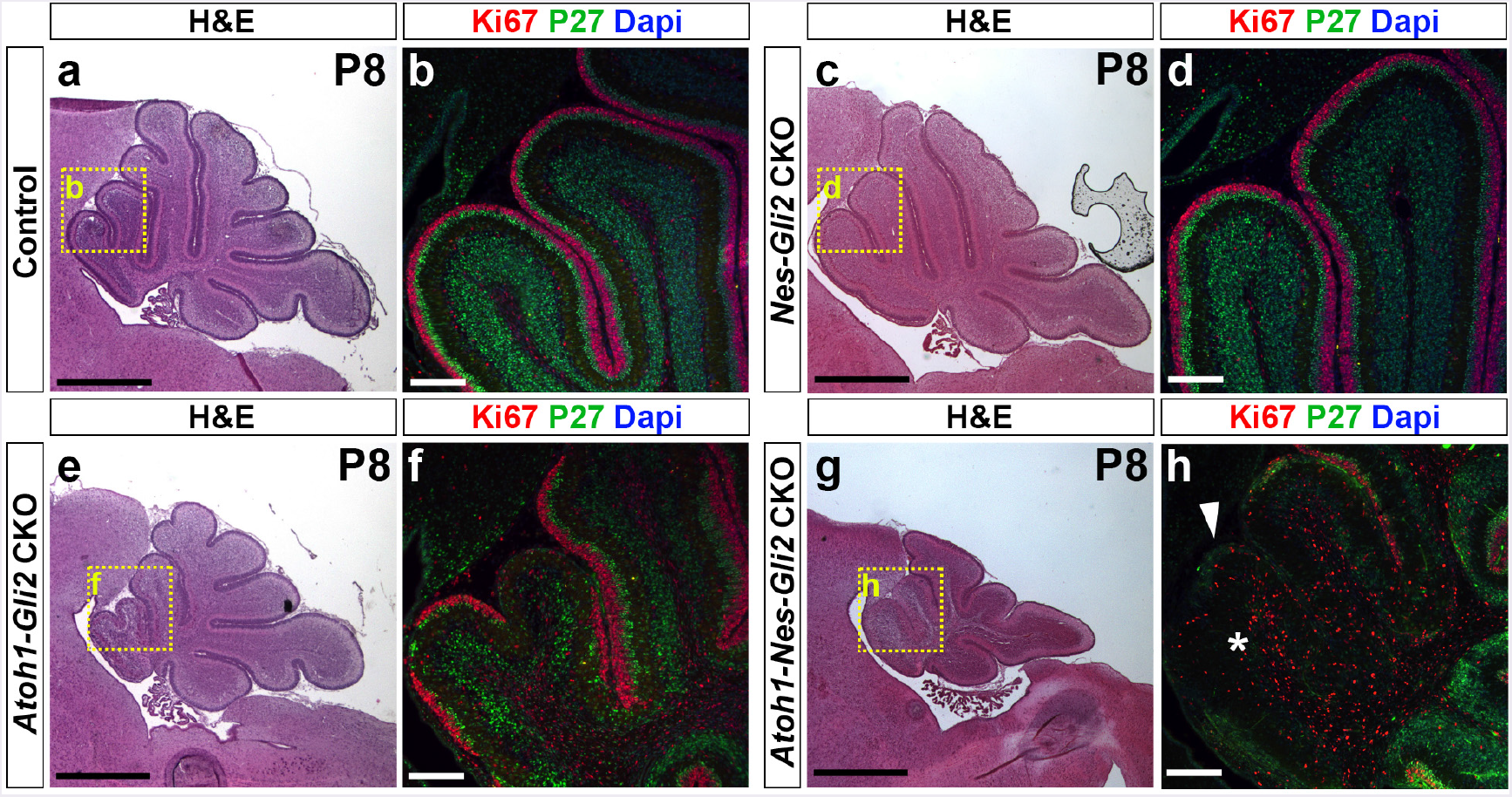
Inactivation of Gli2 in both *Nestin* and *Atoh1* expressing cells inhibits the recovery of the CB. (**a, c, e and g**) H&E of sagittal sections of the cerebellar vermis of P8 *Gli2^flox/flox^* (Control, **a**), *Nes-FlpoER*/+; *R26*^*MASTR*/+^; *Gli2^flox/flox^* (*Nes-Gli2* CKO, **c**), *Atoh1-Cre*/+; *Gli^flox/flox^* (*Atoh1-Gli2* CKO, **e**), and *Atoh1-Cre*/+; *Nes-FlpoER*/+; *R26*^MASTR/^+; *Gli^flox/flox^* (*Atoh1-Nes-Gli2* CKO, **g**) mice injected with Tm at P0. Note that inactivation of *Gli2* only in *Nestin*-expressing cells has no major effect at P8. However, inactivation of *Gli2* in *Nestin*-expressing cells inhibits the compensation mechanism (**g** compared to **e**). (**b, d, f and h**) Close-up (as shown by yellow squares in **a, c, e and g**) of anterior vermis of P8 *Gli^flox/flox^* (**b**), *Nes-FlpoER*/+; *R26*^*MASTR*/+^; *Gli^flox/flox^* (*Nes-Gli2* CKO, **d**), *Atoh1-Cre*/+; *Gli^flox/flox^* (*Atoh1-Gli2* CKO, **f**), and *Atoh1-Cre*/+; *Nes-FlpoER*/+; *R26*^*MASTR*/+^; *Gli^flox/flox^* (*Atoh1-Nes-Gli2* CKO, **h**) cerebella stained with indicated proteins and dapi. White arrowhead and white asterisk indicate the loss of EGL and IGL respectively in *Atoh1-Nes-Gli2* CKO. Scale bars represent 1mm (**a, c, e and g**) and 100μm (**b, d, f and h**).

**Fig. S7.**
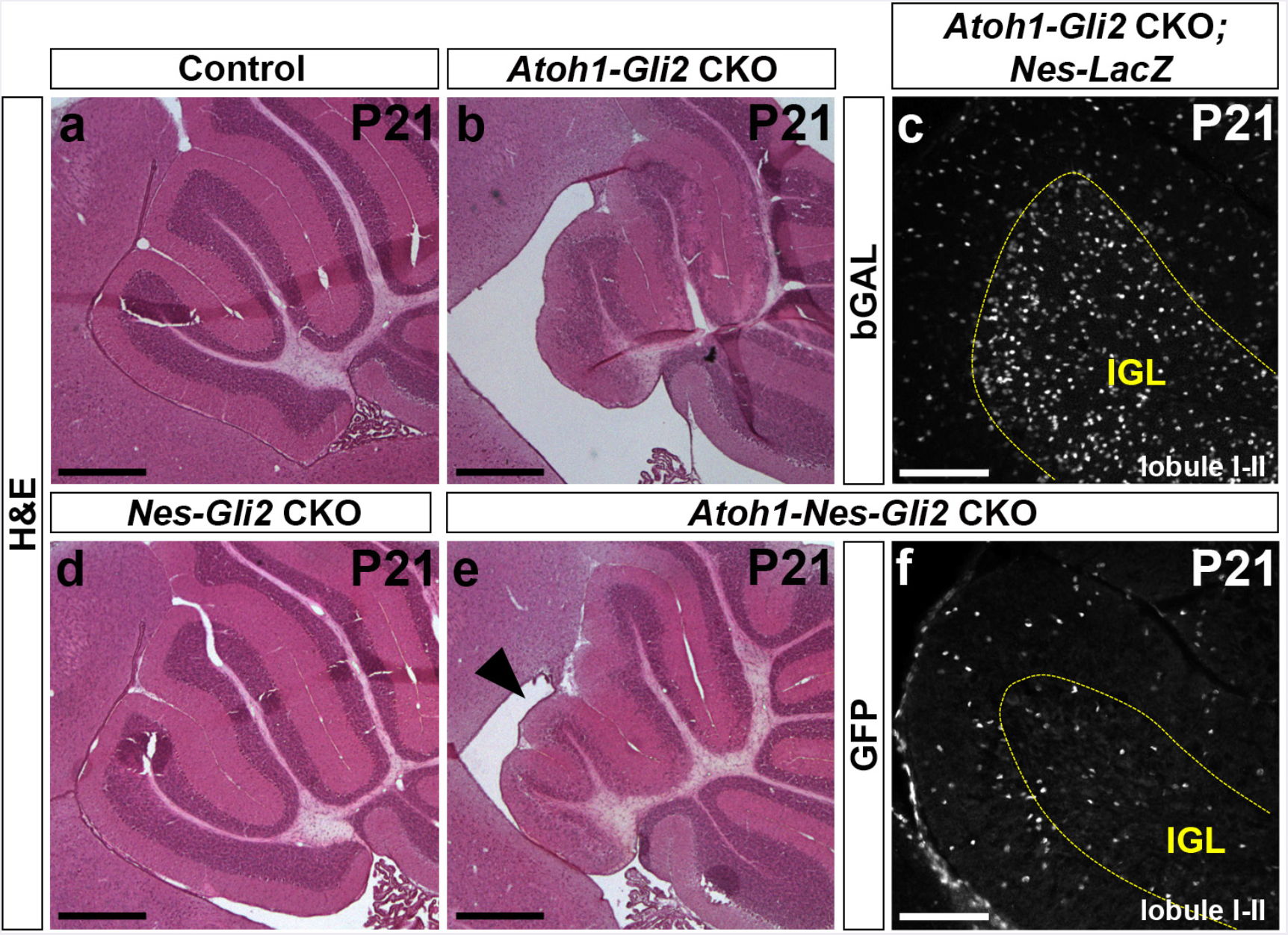
NEPs derived cells failed to populate the IGL at P21. (**a, b, d and e**) H&E of sagittal sections of anterior cerebellar vermis of P21 *Gli2^flox/flox^* (Control, **a**), *Atoh1-Cre*/+; *Gli^flox/flox^* (*Atoh1-Gli2* CKO, **b**), *Nes-FlpoER/+; R26^mastR^+; Gli2^flox/flox^ (Nes-Gli2* CKO, d) and *Atoh1-Cre*/+; *Nes-FlpoER*/+; *R26*^*MASTR*+^; *Gli2^flox/flox^* (*Atoh1-Nes-Gli2* CKO, **e**) mice injected with Tm at P0. Note that inactivation of *Gli2* in Nestin-expressing cells inhibits the compensation mechanism in the anterior vermis (black arrowhead in e). (**c and f**) FIHC detection of the indicated proteins on mid-sagittal cerebellar sections (lobule I-II) of P21 *Atoh1-Cre*/+; *Gli^flox/flox^*; *Nes-FlpoER*/+; *R26*^*FSF-LacZ*/+^ (*Atoh1-Gli2* CKO; *Nes-LacZ*, **c**) and *Atoh1-Cre*/+; *Nes-FlpoER*/+; *R26*^*MASTR*+^; *Gli2^flox/flox^* (*Atoh1-Nes-Gli2* CKO, **f**) mice injected with Tm at P0. IGL is indicated by the yellow doted line. Scale bars represent 500μm (**a, b, d and e**) and 100μm (**c and f**).

## Sup. video1

**P8 WT cerebellum shows no obvious movement of NEPs towards the EGL**. Detection of native CFP fluorescence on sagittal slices of the vermis (lobule 2/3) of P8 *Nes-CFP*/+ mice showing displacement of CFP+ cells. Image stacks were acquired every 5min for 4h.

## Sup. video2

**PCL NEPs migrate toward the EGL in *Atoh1-Gli2* CKO CB at P8**. Detection of native CFP fluorescence on sagittal slices of the vermis (lobule 2/3) of P8 *Atoh1-Cre*/+; *Gli2^flox/flox^*; *Nes-CFP*/+ (*Atoh1-Gli2* CKO; *Nes-CFP*) mice showing displacement of CFP+ cells. Image stacks were acquired every 5min for 4h.

## Sup. video3

**A subset of NEP derived cells migrate from the EGL towards the IGL in *Atoh1-Gli2* CKO CB at P8**. Detection of native CFP fluorescence on sagittal slices of the vermis (lobule 1/2) of P8 *Atoh1-Cre*/+; *Gli2^flox/flox^*; *Nes-CFP*/+ (*Atoh1-Gli2* CKO; *Nes-CFP*) mice showing displacement of CFP+ cells. Image stacks were acquired every 5min for 4h.

## Abbreviations

CB: Cerebellum
SHH: Sonic Hedgehog
EGL: External Granule Layer
GCP: Granule Cell Precursor
NEP: Nestin-expressing progenitor
MASTR: mosaic mutant analysis with spatial and temporal control of recombination
GIFM: Genetic Inducible Fate Mapping
VZ: ventricular zone
PC: Purkinje cell
IGL: internal granule cell layer
HH: Hedgehog
*Ihh:*: Indian hedgehog
*Dhh:*: Desert hedgehog
PTCH1: Patched1
SMO: Smoothened
Ci: cubitus interruptus
A: Activator
R: Repressor
CKO: conditional knockout
P: Postnatal day
Tm: Tamoxifen
IHC: immunohistochemistry
GFP: Green Fluorescent Protein
het: Heterozygous
TUNEL: Terminal deoxynucleotidyl transferase dUTP nick end labeling
PCL: Purkinje cell layer
ISH: *in situ* hybridization
WT: Wild type
TDTom: Tandem Dimeric derivative of DsRed
ML: Molecular layer

